# Graph perceiver network for lung tumor and bronchial premalignant lesion stratification from histopathology

**DOI:** 10.1101/2023.10.16.562550

**Authors:** Rushin H. Gindra, Yi Zheng, Emily J. Green, Mary E. Reid, Sarah A. Mazzilli, Daniel T. Merrick, Eric J. Burks, Vijaya B. Kolachalama, Jennifer E. Beane

## Abstract

Bronchial premalignant lesions (PMLs) precede the development of invasive lung squamous carcinoma (LUSC), posing a significant challenge in distinguishing those likely to advance to LUSC from those that might regress without intervention. In this context, we present a novel computational approach, the Graph Perceiver Network (GRAPE-Net), leveraging hematoxylin and eosin (H&E) stained whole slide images (WSIs) to stratify endobronchial biopsies of PMLs across a spectrum from normal to tumor lung tissues. GRAPE-Net outperforms existing frameworks in classification accuracy predicting LUSC, lung adenocarcinoma (LUAD), and non-tumor (normal) lung tissue on The Cancer Genome Atlas (TCGA) and Clinical Proteomic Tumor Analysis Consortium (CPTAC) datasets containing lung resection tissues while efficiently generating pathologist-aligned, class-specific heatmaps. The network was further tested using endobronchial biopsies from two data cohorts, containing normal to carcinoma in situ histology, and it demonstrated a unique capability to differentiate carcinoma in situ lung squamous PMLs based on their progression status to invasive carcinoma. The network may have utility in stratifying PMLs for chemoprevention trials or more aggressive follow-up.

## Introduction

Lung squamous cell carcinoma (LUSC), the second most common type of non-small cell lung cancer (NSCLC), is preceded by the development of bronchial premalignant lesions (PMLs) that can progress toward invasive carcinoma or regress without intervention. Recently, genomic and proteomic profiling of PMLs has demonstrated that PML progression is associated with impaired immunosurveillance^1–5^. Currently, there are ongoing efforts to build a Lung Pre-Cancer Atlas^6^ to understand molecular alterations in their spatial context associated with disease severity and progression. To date, H&E-stained tissue slides of PMLs have not been fully utilized as a data source to augment our insights into PML biology. Endobronchial biopsies of PMLs are heterogeneous and contain a range of histologic grades, a variety of structures including cartilage and submucosal glands, and varying degrees of inflammatory cell infiltrate. We hypothesized that computational methods that utilize digitized H&E whole slide images (WSIs) may be able to capture PML heterogeneity and stratify PMLs by histologic severity or their ability to progress to invasive carcinoma, providing a standardized and informative WSI-level assessment of PMLs.

We developed a Graph Perceiver Network (GRAPE-Net) to characterize PMLs using lung tissue WSIs from public cohorts (TCGA^7^ and CPTAC^8^ consisting of Normal (non-tumor adjacent), lung adenocarcinoma (LUAD) and LUSC tissue samples. GRAPE-Net was trained using TCGA samples and tested using CPTAC samples. It was then used to learn WSI-level features and classes of endobronchial biopsy-based WSIs from two datasets from the University College London (UCL) and the Roswell Park Comprehensive Cancer Institute (Roswell). The network identified signatures that differentiated the lung tissue types, stratified samples by their predominant histologic pattern within LUSC and LUAD tumors and identified carcinoma in situ (CIS) PMLs that progress to LUSC.

## Methods

### Ethics statement

The Institutional Review Boards at Boston University’s Chobanian and Avedisian School of Medicine and the Roswell Park Comprehensive Cancer Center approved the study. All subjects provided written informed consent. Data from TCGA, CPTAC, and UCL are publicly available.

### Data acquisition and preprocessing

We obtained digitized H&E images of lung resection tissue and lung tumor subtype annotation from TCGA (534 Normal, 740 LUSC, 808 LUAD) and CPTAC (719 Normal, 685 LUSC, 667 LUAD)^8^. We also obtained the predominant tumor histologic pattern for the CPTAC samples. The endobronchial biopsy-based WSIs were from two datasets: 1) University College London (UCL) (112 CIS samples)^4^, defining progression as moving to LUSC; and 2) Roswell Park Comprehensive Cancer Institute (Roswell) (346 samples, normal to CIS spectrum)^1^, where progression was persistence of dysplasia or advancement to at least mild dysplasia (**Figure 1a**). The UCL CIS samples included supplementary details about the samples’ lymphocyte counts in CIS and stromal regions. The Roswell samples included molecular phenotypes identified using RNA sequencing data such as PML molecular subtype that included 4 categories – proliferative, inflammatory, secretory, and normal-like, as previously described^1^.

**Figure 1:**
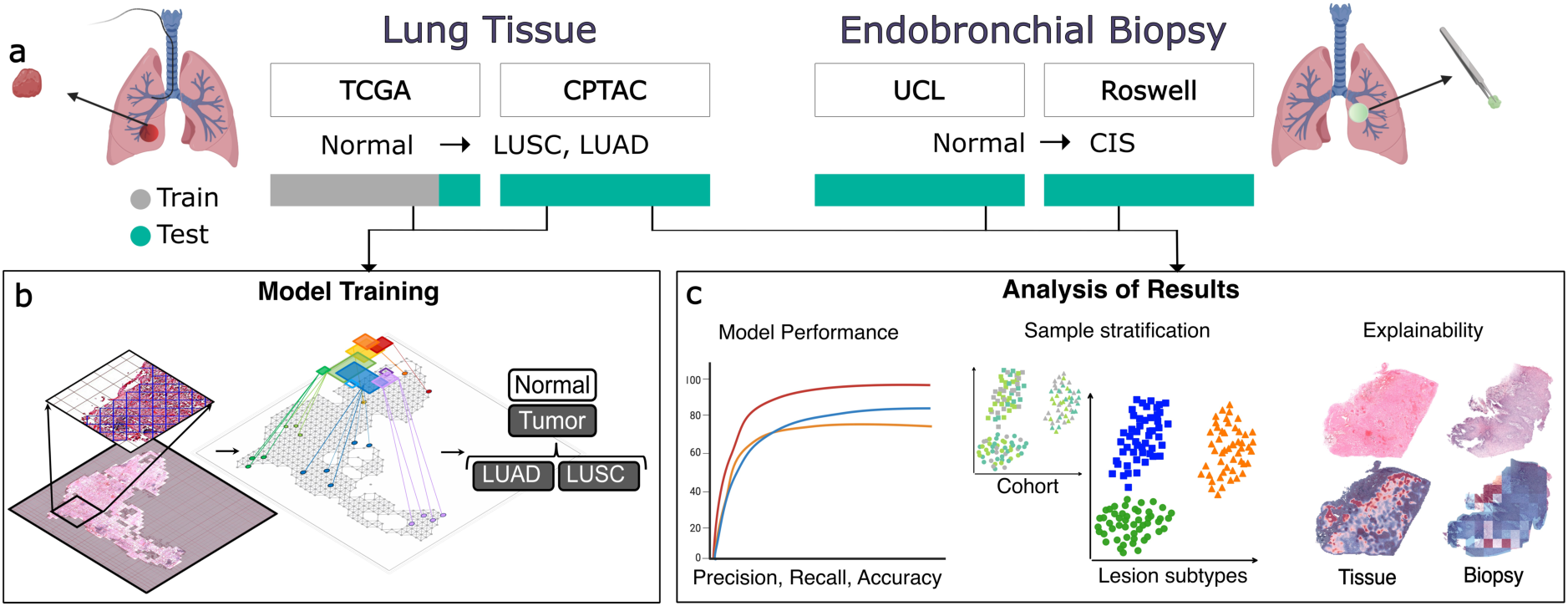
Overview of the study. **(a)** Digitized H&E-stained images were obtained from 4 different datasets including lung resection tissues (TCGA and CPTAC datasets) with normal tissue adjacent to the tumor (Normal), lung adenocarcinoma (LUAD), and lung squamous cell carcinoma (LUSC) and endobronchial biopsies (UCL and ROSWELL datasets) ranging from normal to carcinoma in situ (CIS) histology. The model was trained on TCGA data and test across TCGA, CPTAC, UCL, and ROSWELL datasets. **(b)** Graph Perceiver Network (GRAPE-Net) overview trained to differentiate lung cancer subtypes. **(c)** Overview of analysis conducted using the results of the model that included evaluation of model performance, WSI features with clustering-based metrics, and model explainability using class-specific heatmaps.

To evaluate our model’s explanations, we selected a few WSIs from CPTAC, Roswell, and UCL and uploaded them to a web-based software (PixelView, deepPath, Boston, MA). Several regions on the WSIs were annotated by board certified thoracic pathologists (D.M. and E.B.) using the software. In the LUAD specimens, tumor areas were annotated by their histologic patterns (solid, micropapillary, cribiform, papillary, acinar, and lepidic). For both LUAD and LUSC tumors, histologic features, including necrosis, lymphatic, and vascular invasions, were also annotated. Airways present in the tissues were graded on the spectrum of normal to CIS and non-tumor tissue areas were labeled as healthy or inflamed lung tissue, stroma, or necrosis. The LUAD and LUSC annotations were combined into tumor, stroma, normal lung, and necrosis for analysis. For the endobronchial biopsy samples, epithelial regions were annotated based on histologic grade (normal, hyperplasia, immature squamous metaplasia, squamous metaplasia, mild dysplasia, moderate dysplasia, severe dysplasia, and CIS). These annotations were combined into CIS, dysplasia, and non-dysplasia for analysis. The overlap between the model-produced heatmaps and the pathologist annotations was analyzed by exporting them as binary images.

WSIs are inherently large, often exceeding dimensions of tens of thousands of pixels in both width and height. This presents challenges for analyzing them due to computational and memory constraints. To address these limitations, each WSI was passed through a fast-patching pipeline at 20x magnification, creating non-overlapping tissue region patches of 256×256 pixels while filtering out background slide information. Key epithelial regions were retained via Otsu thresholding^9^. To prepare the data as input for GRAPE-Net, each WSI was represented as an undirected, unweighted graph with nodes representing the tissue image patches and set of edges connecting the nodes. The graph follows an 8-connectivity neighborhood structure. The node embeddings in the graph were structured as a matrix of *N* feature vectors, where *N* is the total number of patches in the WSI (nodes in the graph). Each vector is of size 768-dimensions. The features for each patch were obtained from CTransPath^10^, a swin-transformer^11^ based feature extractor. CTransPath was pre-trained on the TCGA pan-cancer dataset in a self-supervised manner. The graph neighborhood connections are structured as a binary *N* × *N* matrix (adjacency matrix *A*) with elements *e_ij_* = 1 if there exists an edge between nodes *i* and *j*. These graphs feed into GRAPE-Net, which was then trained for a WSI classification task (**Figure 1b**).

### Graph perceiver network

The architectural design of GRAPE-Net consists of three main components (**Figure 2**). (1) A graph convolution block^12^ that retains positions information in the input features via edge connections between neighboring nodes, (2) a cross-attention block (PMA) from perceiver^13^ which maps variable number of patches per WSI graph to predefined clusters (or “sets”), (3) a self-attention block (SAB)^14^ that learns relevant interactions between the sets.

**Figure 2:**
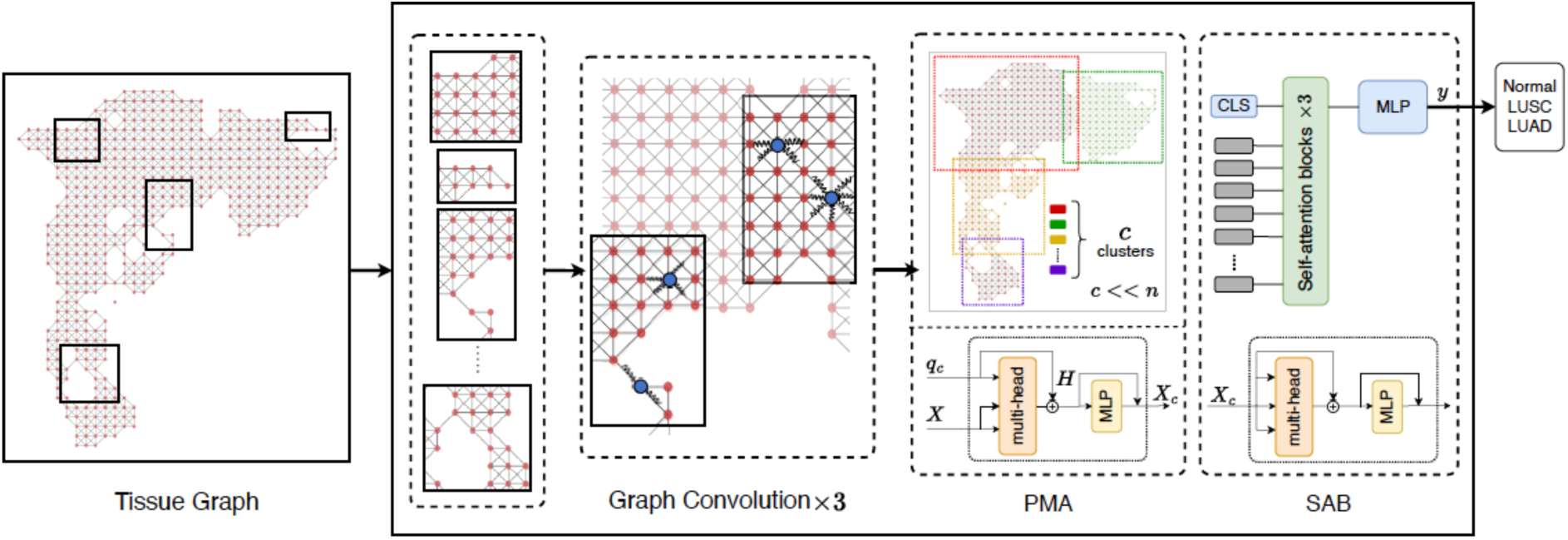
Graph perceiver network (GRAPE-Net) architecture. Each whole slide image (WSI) is represented as an undirected, unweighted tissue graph, where each node is an embedding of features in an image patch. Our proposed GRAPE-Net aggregates neighborhood information via graph convolutions while preserving spatial context^12^. To efficiently find the neighborhood interaction between the different tissue regions in the tumor micro-environment, the graph is clustered to ‘*C*’ overlapping sets by pooling the nodes using multi-head cross-attention block (PMA). The fixed set embeddings are then given as input to the self-attention block (SAB), which learn morphologic interactions with its neighborhood and relevance of each set towards the specific output label of interest. The aggregated attentions are then given as input to a multi-layer perceptron (MLP) for classification of the WSI as either Normal, LUAD or LUSC. Here, the layer-specific feature representation of the graph is denoted as H. A CLS (or class) token is added to serve as the entire WSI representation, which is used for classification and computing the relevance of each graph node towards the prediction.

The goal of GRAPE-Net is to learn a representation of the WSI. The graph convolution block contributes to this by stacking multiple graph convolutional layers to learn hierarchical features efficiently. Specifically, the first few layers capture the local, low-level patterns while the deeper layers aggregate them to understand the global structures within the graph. Next, inspired by the recent work on using min-cut pooling on a graph-transformer network (GTP) for WSI level classification^15^, and adaptive aggregation functions used in recent graph classification problems^16^, GRAPE-Net uses a cross-attention block (PMA) from the perceiver^13^. This module simplifies GTP by replacing the min-cut pooling^17^ mechanism with cross-attention layers, which cluster the graph nodes, with similar embeddings and spatial proximity, to predefined latent clusters which we denote as “sets”. The number of sets used in the PMA module is a hyperparameter *C*, typically much smaller than the expected number of visual tokens. The formed sets, accompanied by a classification token CLS, are inputs to the self-attention block, mirroring the approach employed in vision transformers^18^. The SAB assigns weights to each set embedding for classification. The PMA and SAB modules are alternatively stacked, allowing the set embeddings to extract information from the input WSI graph based on the previous layer’s learnings. This strategy preserves positional information of the graph nodes in the sets. We followed the weight sharing strategy of the perceiver to optimize computations and reduce memory constraints. The final SAB block is followed by a multi-layer perceptron (MLP), which facilitates multi-class prediction. The CLS token embedding from the SAB represents the slide-level aggregation vector of the weighted sets. The utilization of the CLS token instead of averaging the embeddings of sets facilitates explainability in the network.

Our model introduces novelty by integrating the graph module with the perceiver architecture, enabling sparse graph computations on the visual tokens and computationally efficient modeling of a graph for downstream tasks. For our use-case, it serves as an end-to-end, amortized clustering algorithm, which clusters distinct regions of the tissue into overlapping sets, preserving vital spatial interactions that enhance our comprehension of the biological processes withing tumor tissues.

### Experimental design

GRAPE-Net was trained as a three-label classifier (LUAD, LUSC, or Normal). We used the TCGA dataset, with stratified patient-level sampling and 5-fold cross-validation (**Supplementary Figure 1**) with internal testing. The receiver operating characteristic (ROC) and precision-recall (PR) curves for each class were computed along with the accuracy, precision, sensitivity, and specificity as performance metrics (**Figure 1c**). To ensure the classifier is robust to dataset specific batch effects, it was evaluated on the unseen CPTAC cohort. Model weights of the fold with the highest sensitivity (recall) on CPTAC cohort was used for further post-hoc analysis.

To evaluate the effectiveness our proposed network, we compared the model performance with a state-of-the art lung tumor classification model, GTP^15^. Since our model leverages graph convolutions, we also selected a traditional graph classifier, Graph Isomorphism Network (GIN)^12^ for comparison. To ensure fair comparison, we used CTransPath^10^ as the common feature extractor for all methods and used the same cross-validation parameters for training. The hyper-parameters stated in the respective published research of GTP and GIN were used to achieve the best performance on TCGA and CPTAC cohorts. Optimal hyperparameters for our network configuration involved a hidden dimension (D) of 64, a graph block with 3 GINConv layers, a cross-attention pooling configuration of 200 sets and a 3 self-attention layers per self-attention block (SAB). We used 8 heads for multi-head attention training. To avoid overfitting, we used dropouts with a rate of 0.2 in feed-forward blocks, and binary cross-entropy loss to enhance robustness. Training was completed over 30 epochs in 8-sample mini-batches, using early stopping, a 0.0003 initial learning rate, a step scheduler, and the Adam optimizer^19^ for expedited convergence. All the experiments were performed on a single GeForce RTX 2080Ti 11Gb workstation.

For post-hoc analysis, the WSI-level features for each sample were extracted by giving the respective graphs as input to the trained 3-label GRAPE-Net classifier. The 64-dimensional features from the final layer of the network were considered as the representations for each WSI and used for further analysis. UMAP manifold representations^20^and principal component analysis (PCA) representations were plotted to illustrate the relationship between image features and sample phenotypes (**Figure 1c**). Sample phenotypes were grouped as follows: normal adjacent lung tissue, LUSC, and LUAD for the lung tissue data (CPTAC). Within CPTAC, we examined differences in the principal components associated with LUSC predominant histologic patterns (keratinizing or non-keratinizing) and LUAD predominant histologic patterns grouped as follows: (solid/micropapillary/cribriform or lepidic/acinar/papillary). Within the CIS samples, we examined differences in the principal components by progression status. Post-hoc analysis results suggest that the model is robust to cohort-based batch effects between Roswell and UCL cohorts.

### Explainability analysis

GRAPE-Net learns the classification label-relevant contributions of the tissue regions during training. We show these explanations as heatmaps using a relevance propagation mechanism^21^(**Figure 1c**). This mechanism is a generic attention-based model explanations for bimodal transformers. It uses attentions and gradients from the SAB and PMA blocks to produce relevancy maps for each interaction between the sets *C* and the assigned nodes *N*. The relevancy map from PMA provides the individual contributions of each node (patch) for the final prediction instead of distributing the contribution of each set as an average to all the nodes within the set. These relevancy maps are constructed on the WSI using the adjacency matrix *A* and coordinates of all the patches on the WSI.

### Statistical analysis

We compared the performance of GRAPE-Net with other approaches for the tumor classification task in lung cancer (train on TCGA-Lung, test on CPTAC-Lung) highlighting its efficiency while performing like the state-of-the-art methods. UMAP and PCA clustering analyses were conducted using Scanpy framework ^22^. Clustering performance was evaluated using adjusted Rand score and adjusted mutual index with higher scores displaying good clustering for user-specified labels. Statistical significance between principal components and sample characteristics were assessed through two-tailed t-tests and Spearman’s correlation coefficient experiments with analyses performed at a significance *p* = 0.05.

### Data availability

Data from TCGA and CPTAC can be downloaded freely from the public domain. UCL data can be obtained from: https://idr.openmicroscopy.org with IDR0082. Roswell images can be accessed via the Human Tumor Atlas Network Data Portal (https://humantumoratlas.org/) using the file identifications provided in **Supplementary Table 1**.

### Code availability

Computer scripts and manuals are made available on GitHub (https://github.com/vkola-lab/ajpa2024).

## Results

### GRAPE-Net Performance

GRAPE-Net demonstrated high classification performance on TCGA (mean AUROC=0.98 ± 0.01 & mean AUPRC=0.98 ± 0.01) and CPTAC data (mean AUROC=0.93 ± 0.01 & mean AUPRC=0.95 ± 0.02) (**Figure 3**), utilizing significantly fewer resources compared to two published frameworks (**Table 1**), *GRAPE-Net* ~ 202K vs *GTP* ~ 664K trainable parameters). Five-fold cross validation indicated good agreement between the true and predicted tumor classes on both the TCGA and CPTAC data, across different folds displaying the robustness of the model to data bias used for training and testing. For each sample, three separate class activation maps were produced to highlight the areas of normal and tumor tissue (LUAD or LUSC) in each WSI. The generated class-specific heatmaps were consistent with the expert pathologist annotations from 20 CPTAC samples (selected at random), accurately pinpointing areas corresponding to same-label annotations (**Figure 4a** and **Supplementary Figure 3**). These results illustrate that GRAPE-Net can distinguish between the NSCLC tumor subtypes and the normal tissue regions within a WSI, and therefore, we wanted to further explore its ability to detect subtle distinction between tumor histologic patterns and PML histology.

**Figure 3:**
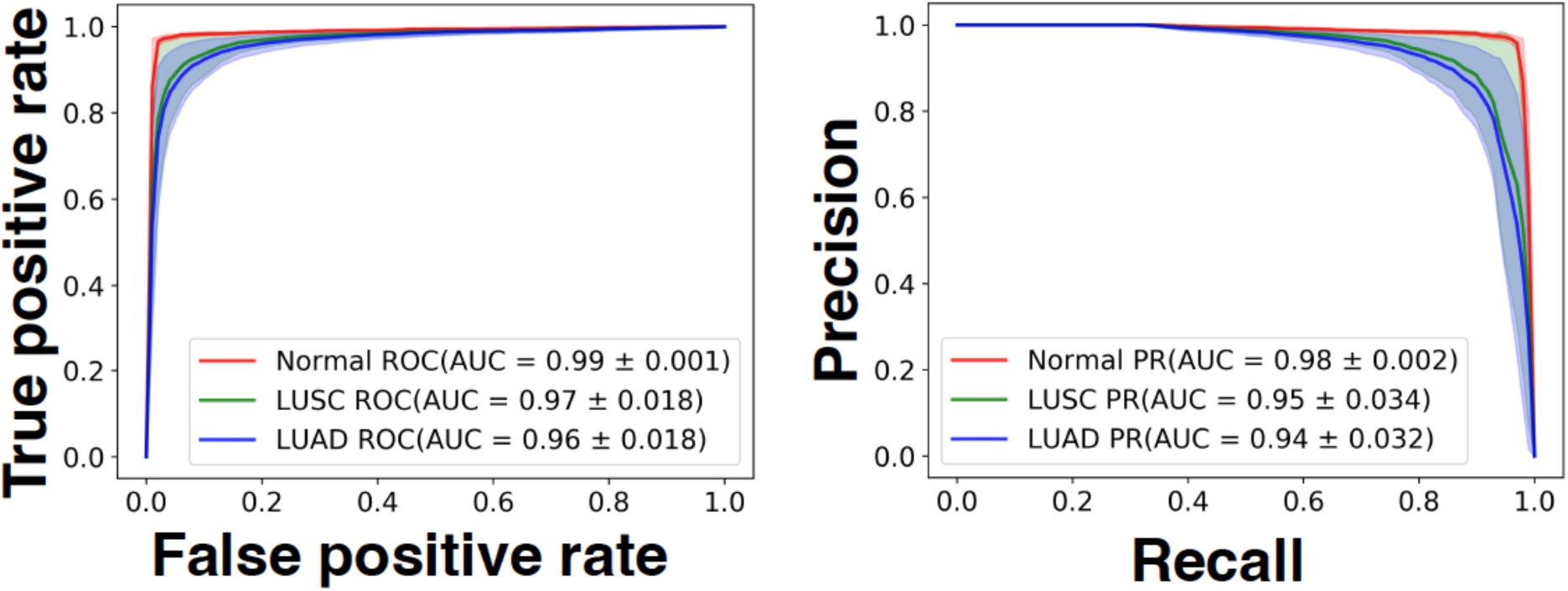
Classification Performance. Receiver Operating Characteristic (ROC) and Precision-Recall (PR) curves showcasing the performance of GRAPE-Net towards multi-class classification on CPTAC external testing dataset. The area under the curve (AUC) score for each label (Normal, LUAD, LUSC) is provided for the ROC and PR curves respectively.

**Figure 4:**
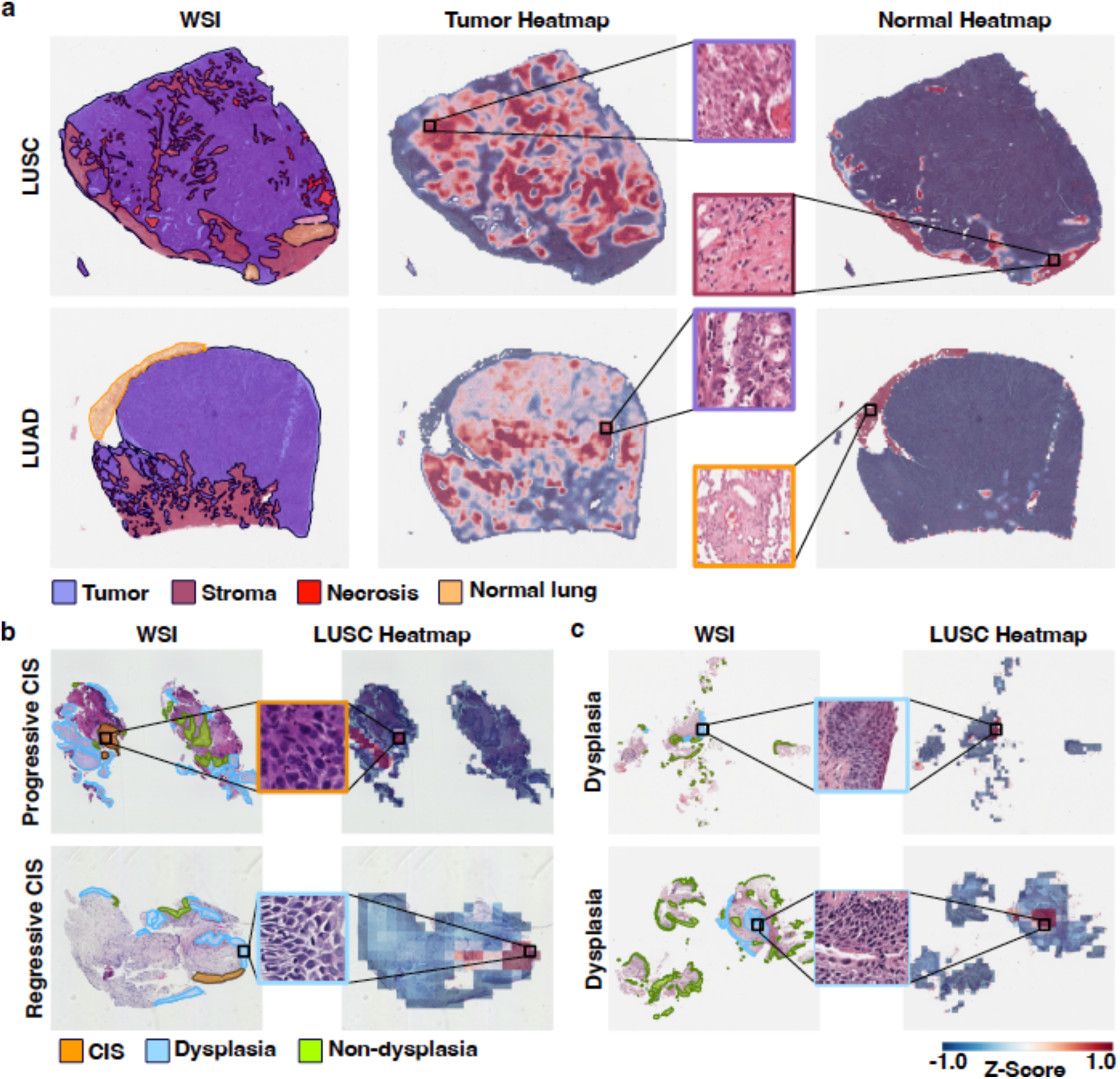
GRAPE-Net identifies tissue regions that correspond with pathologic annotations. **(a)** The network identified regions annotated as tumor and normal on LUAD and LUSC WSIs from CPTAC. **(b, c)** The network was also used to classify PMLs as one of the three labels. The LUSC-specific heatmaps for PMLs predicted to be LUSC identified regions of dysplasia and CIS as well as other regions that are likely contributing to its ability to separate CIS progressors from CIS regressors.

**Table 1:**
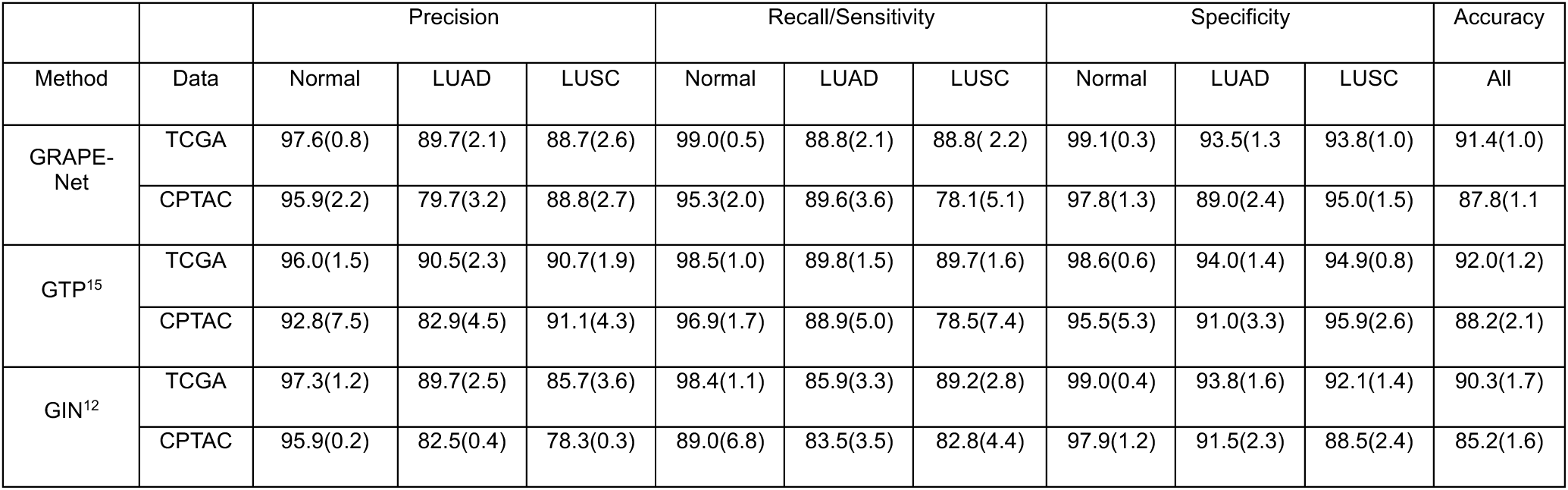
Performance metrics for the 3-label (Normal vs LUAD vs LUSC) classification task. Mean performance metrics(%) with standard deviation values in parentheses are reported for the GRAPE-Net method used in this study and two other previously published state of the art methods in the TCGA and CPTAC datasets.

### Stratification of Tissues by Morphologic Features

We hypothesized that GRAPE-Net could differentiate between predominant histologic patterns of LUSC and LUAD tumors and lung squamous PML histology by discerning subtle morphological features. To test this, we used the WSI latent features from the CPTAC, UCL, and Roswell samples to perform UMAP-based clustering (**Figure 5**). Clustering segregated normal, LUAD, and LUSC with a high adjusted Rand score of 0.69 and adjusted mutual information score of 0.64. Principal component analysis (PCA) on the latent features (**Supplementary Figure 2**) revealed significant PC1 differences between keratinizing and non-keratinizing LUSC tumors (PC1: *p* = 0.01), with the latter PC2 values like those of LUAD tumors. The LUAD tumors showed significant PC1 differences between aggressive (solid/micropapillary) and non-aggressive (lepidic/acinar/papillary) histologic patterns (PC1: *p* = 5.4*e-*14). Notably, LUAD (solid/micropapillary) tumors exhibited PC2 values similar to LUSC tumors, as solid LUAD tumors require immunohistochemical analysis for diagnosis as they can be difficult to distinguish from LUSC tumors.^23^

**Figure 5:**
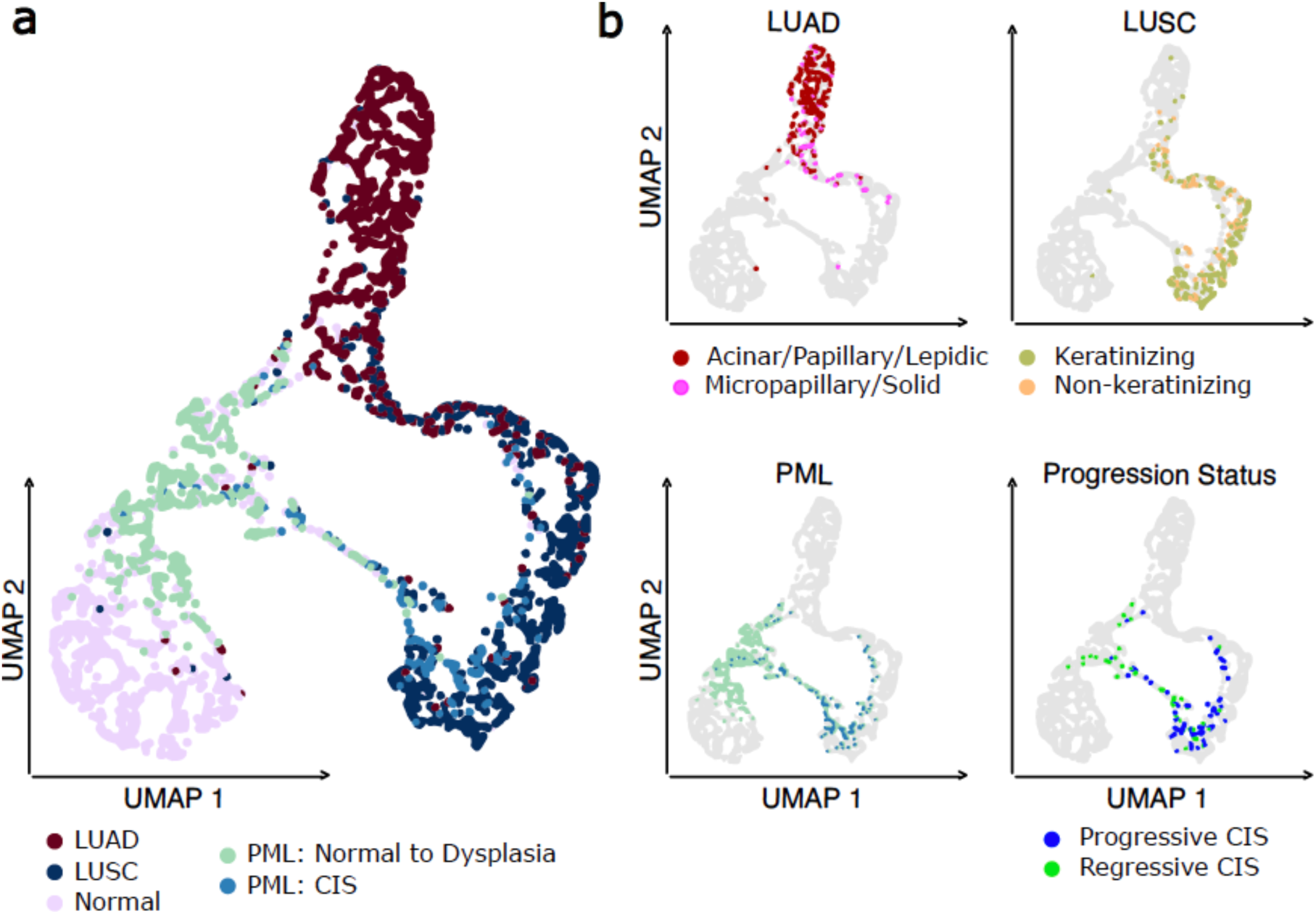
Stratification of lung tumors and PMLs relates to histologic features and outcome. UMAP plots of CPTAC, UCL, and Roswell WSI features stratified samples from normal to invasive carcinoma. The model separated LUAD and LUSC WSIs by tumor histologic patterns and CIS PMLs by outcome.

UMAP and PCA revealed most endobronchial biopsies were proximal to the normal cluster, with some high-grade dysplasia and CIS PMLs clustering with LUAD and LUSC cases (*p <* 0.01 for PML vs. LUAD, PML vs. LUSC, on both PC1 and PC2). Roswell biopsies did not show significant separation by histology, molecular subtype, or progression status (**Supplementary Figure 2c**).

All CIS samples from Roswell and some from UCL grouped with LUSC tumor cases (75% of CIS samples were classified as LUSC tumors with *p <* 0.01 for CIS vs. LUAD, CIS vs. LUSC on both PC1 and PC2). Among the UCL biopsies with progression status to LUSC, the model robustly stratified regressive versus progressive CIS. GRAPE-Net performance predicting regressive CIS as Normals (pNORMAL) and Progressive CIS as Tumors (pLUSC) was as follows: Precision = 0.68, Recall = 0.67, Specificity = 0.38, Accuracy = 0.67, and AUC=0.69 (**Table 2**). Additionally, in prior work on the UCL dataset^4^, an automated deep learning pipeline was used to quantify the density of lymphocytes, epithelial cells, and stromal cells in the CIS and stromal regions. The study showed a significant decrease in lymphocyte density (p=0.02) and increase in epithelial cell density (p=0.005) in CIS regions in progressive versus regressive samples. Using GRAPE-Net, we observed a similar significant decrease in lymphocyte density (p=1.03e-6) and an increase in epithelial cell density (p=1.4e-4) between CIS samples predicted as Tumors (pLUSC) versus Normals (pNORMAL). Spearman correlation between the PC values and the respective CIS region lymphocyte and epithelial cell densities suggested that PC1 has a high positive correlation and PC2 has a high negative correlation with epithelial density. PC2 is also positively correlated with lymphocyte density (**Table 2**). Both PC1 and PC2 are significantly different between CIS progressive versus regressive lesions (p=0.005 and p=1.1e-5, respectively). These results suggest that the model learns certain morphological characteristics pertaining to CIS progression status highlighted using relevance heatmaps as shown in **Figure 4b** and **Supplementary Figure 4**.

**Table 2:** Significance statistics for comparing model feature space with sample characteristics for CIS premalignant lesions. To demonstrate GRAPE-Net’s ability to stratify PMLs, we show the model’s classification performance predicting regressive CIS as Normals (pNORMAL) and Progressive CIS as Tumors (pLUSC). Additional test statistics were also done with other metadata characteristics, with p-value<0.05 considered significant. Significant scores highlighted in **bold.**

| Test Description | Test Statistic | P-Value (<0.05) |
| --- | --- | --- |
| **T Tests** |  |  |
| Lymphocyte Density in CIS regions (pLUSC v pNORMAL) | **-5.18** | **1.03e-06** |
| Epithelial Density in CIS regions (pLUSC v pNORMAL) | **3.94** | **0.000145** |
| PC1 (Progression v Regression) | **2.82** | **0.005721** |
| PC2 (Progression v Regression) | **-4.59** | **1.18e-05** |
| **Spearman Correlation Coefficient** |  |  |
| Lymphocyte Density in CIS regions v PC1 | -0.099 | 0.2932 |
| Lymphocyte Density in CIS regions v PC2 | **0.297** | **0.0024** |
| Epithelial Density in CIS regions v PC1 | **0.506** | **0.0002** |
| Epithelial Density in CIS regions v PC2 | **-0.749** | **0.0002** |
| **Classification Metrics** |  |  |
| Precision | 0.68 | - |
| Recall | 0.67 | - |
| Specificity | 0.38 | - |
| Accuracy | 0.67 | - |
| AUC | 0.69 | - |

## Discussion

In this work, we introduce the Graph Perceiver Network (GRAPE-Net) that is designed to stratify bronchial PMLs which are precursors to invasive LUSC. The goal was to distinguish PMLs with a high likelihood of progressing to LUSC from those that may regress without intervention. Utilizing H&E stained WSIs, GRAPE-Net stratifies PMLs across a continuum from normal to tumor tissues. Using four distinct datasets, the model generated latent features observable across sample types (resected lung tissue versus endobronchial biopsies) that were associated with tumor histologic patterns and CIS PMLs progression status to invasive carcinoma.

GRAPE-Net’s performance was demonstrated through its high classification accuracy on TCGA and CPTAC lung resection tissue datasets, coupled with its efficient generation of pathologist-aligned, class-specific heatmaps. Principal component analysis (PCA) on the WSI latent features demonstrated separation within LUAD and LUSC samples based on predominant histologic pattern. On the endobronchial biopsy WSIs, GRAPE-Net identified CIS lesions that progressed to LUSC. The principal components of the latent features were associated with decreases and increases in lymphocyte and epithelial cell densities within CIS regions, respectively, in progressing versus regressing lesions. GRAPE-Net’s predictive capacity on CIS lesions may have clinical utility in prioritizing patients for chemoprevention trials or by potentially shortening screening intervals via bronchoscopy.

Our framework currently does not differentiate between biopsies with and without bronchial dysplasia, likely due to a paucity of training data on the cellular morphology changes characteristic of various dysplasia stages (i.e., mild, moderate, and severe). This limitation underscores the necessity for future research to expand model training with a broader array of bronchial PMLs. Additionally, it will be important to test our model on lung adenomatous premalignant lesions (including atypic adenomatous hyperplasia and adenocarcinoma in situ) as biopsies of these lesions become available via robotic bronchoscopy. Looking forward, we advocate for the assessment of GRAPE-Net across more extensive cohorts and the integration of additional bulk, single cell, and spatial resolved molecular data into the model to refine PML stratification.

## Acknowledgment

We would like to thank Ms. Erin Kane for coordinating the upload of the Roswell slides to the Human Tumor Atlas Network public data portal.

## Author Contributions

Study conception (JEB, VBK); method development (RHG, YZ, VBK, JEB); interpretation of results (RHG, VBK, JEB, EJB); pathologic annotation (EJB, DTM, EJG); sample acquisition and clinical data curation (MER, SAM, JEB); writing of the manuscript (RHG, VBK, JEB).

---

**Supplementary Figure 1:**
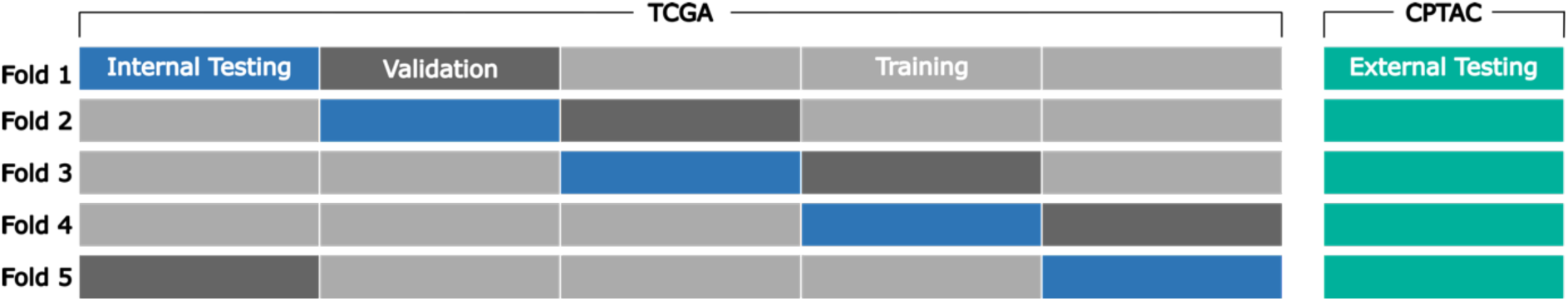
Cross validation protocol. In addition to following the 5 fold cross-validation protocol for training, we also cycle through an unseen internal testing set every fold, to get an unbiased perspective of the model performance. Following the cross-validation, the model is also then tested on an external test set.

**Supplementary Figure 2:**
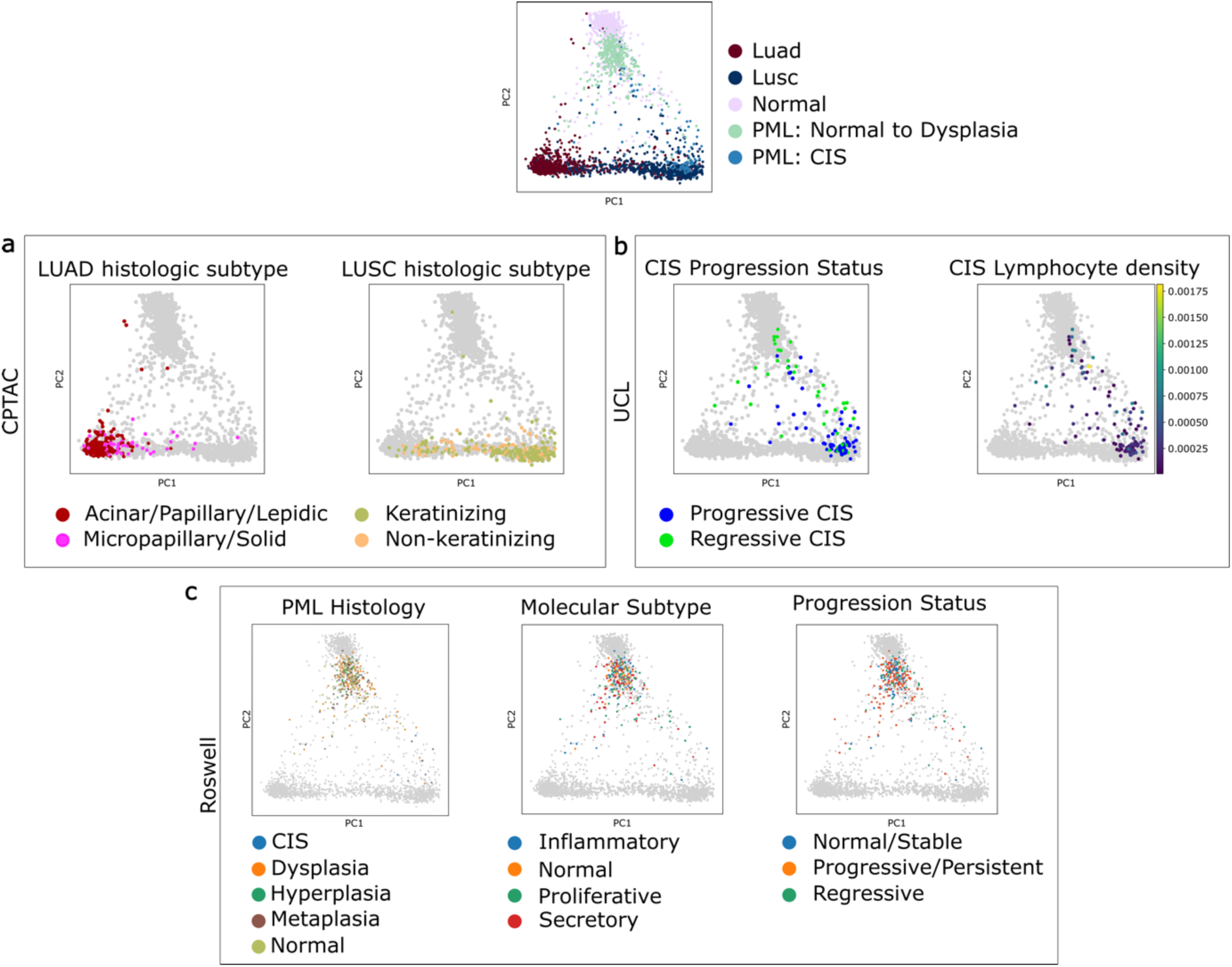
Principal component analysis of lung tumor and premalignant lesions samples. Scatterplots of the first 2 principal components calculated using the final model-derived features across the same samples used for UMAP analysis in Figure 1. (a) Principal component analysis (PCA) plots with CPTAC samples highlighted by tumor subtype (b) PCA plots with UCL samples highlighted by progression status and lymphocyte count (c) PCA plots with Roswell samples highlighted by histologic grade, molecular subtype, and progression status.

**Supplementary Figure 3:**
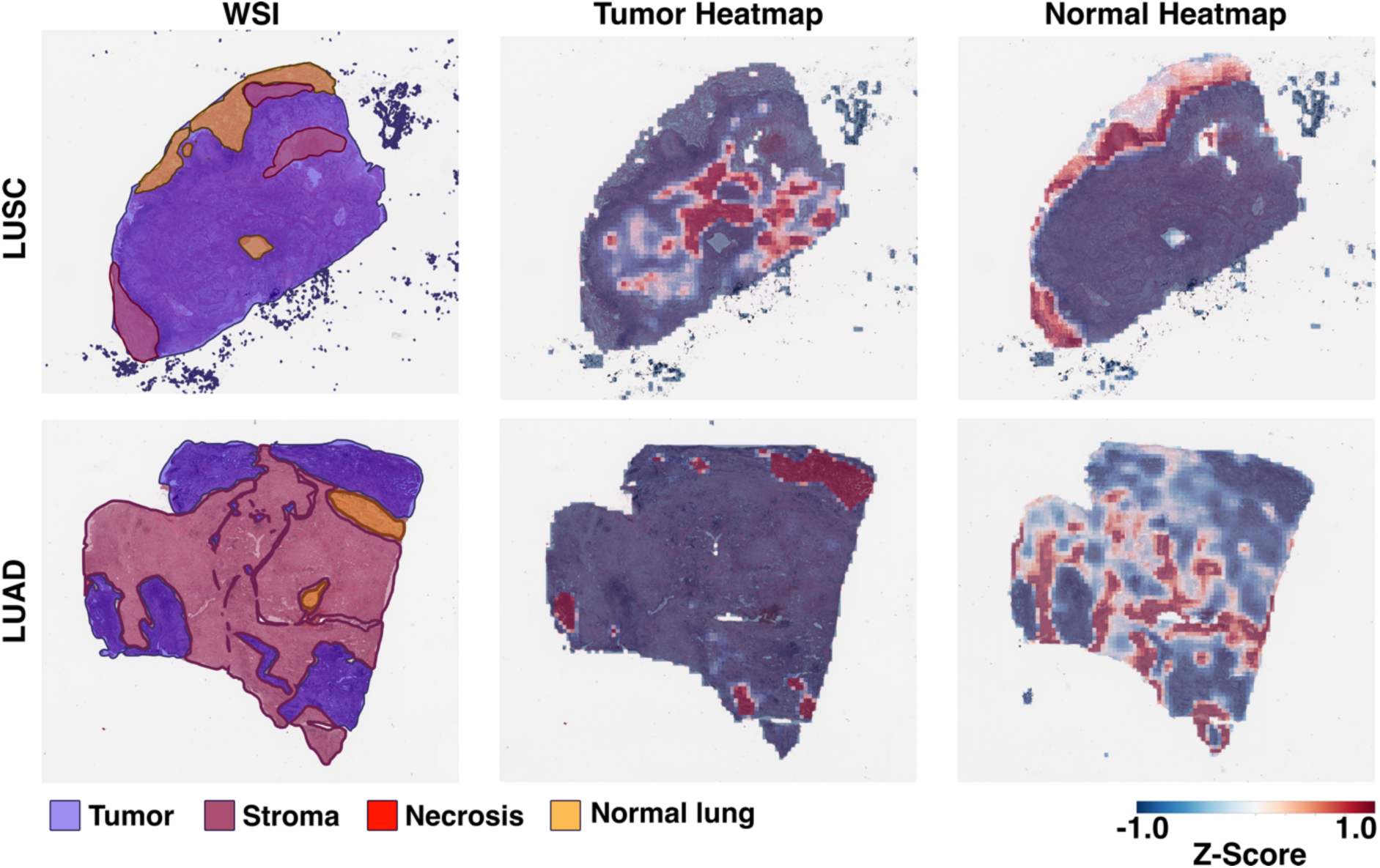
Representative heatmaps of a LUAD and a LUSC case from CPTAC. The procedure for explanation visualizations is identical to Figure 2a. WSIs, their overlayed pathologist annotations (1st column) and respective tumor-specific (2nd column) and normal-specific heatmaps (3rd column) are shown for tumor resection samples from CPTAC cohort (LUSC, upper row and LUAD, bottom row). We observe a strong overlap between the tumor annotations and the tumor-specific heatmaps. Likewise, there is a strong overlap between stromal, necrotic and normal lung annotations and normal-specific heatmaps from GRAPE-Net, showing the network is able to understand the tissue region morphology well.

**Supplementary Figure 4:**
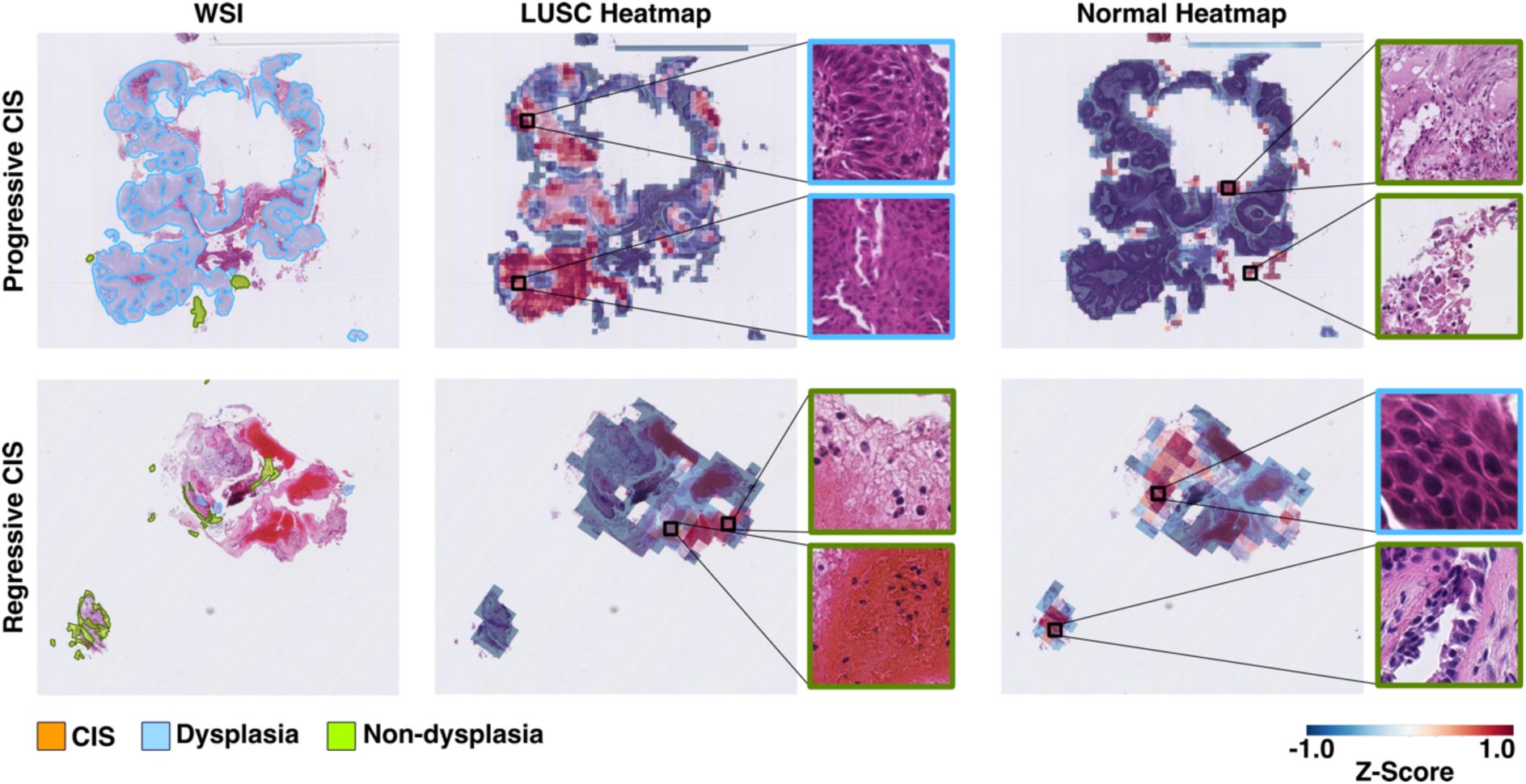
Additional examples of heatmaps generated for UCL samples. The procedure for explanation visualizations is identical to Figure 2b. WSIs, their overlayed pathologist annotations (1st column) and respective LUSC-specific (2nd column) and normal-specific heatmaps (3rd column) are shown for endobronchial premalignant biopsies from UCL. We observe a strong overlap between the dysplasia annotations and LUSC-specific heatmaps in progressive cases but not in regressive cases.

**Supplementary Figure 5:**
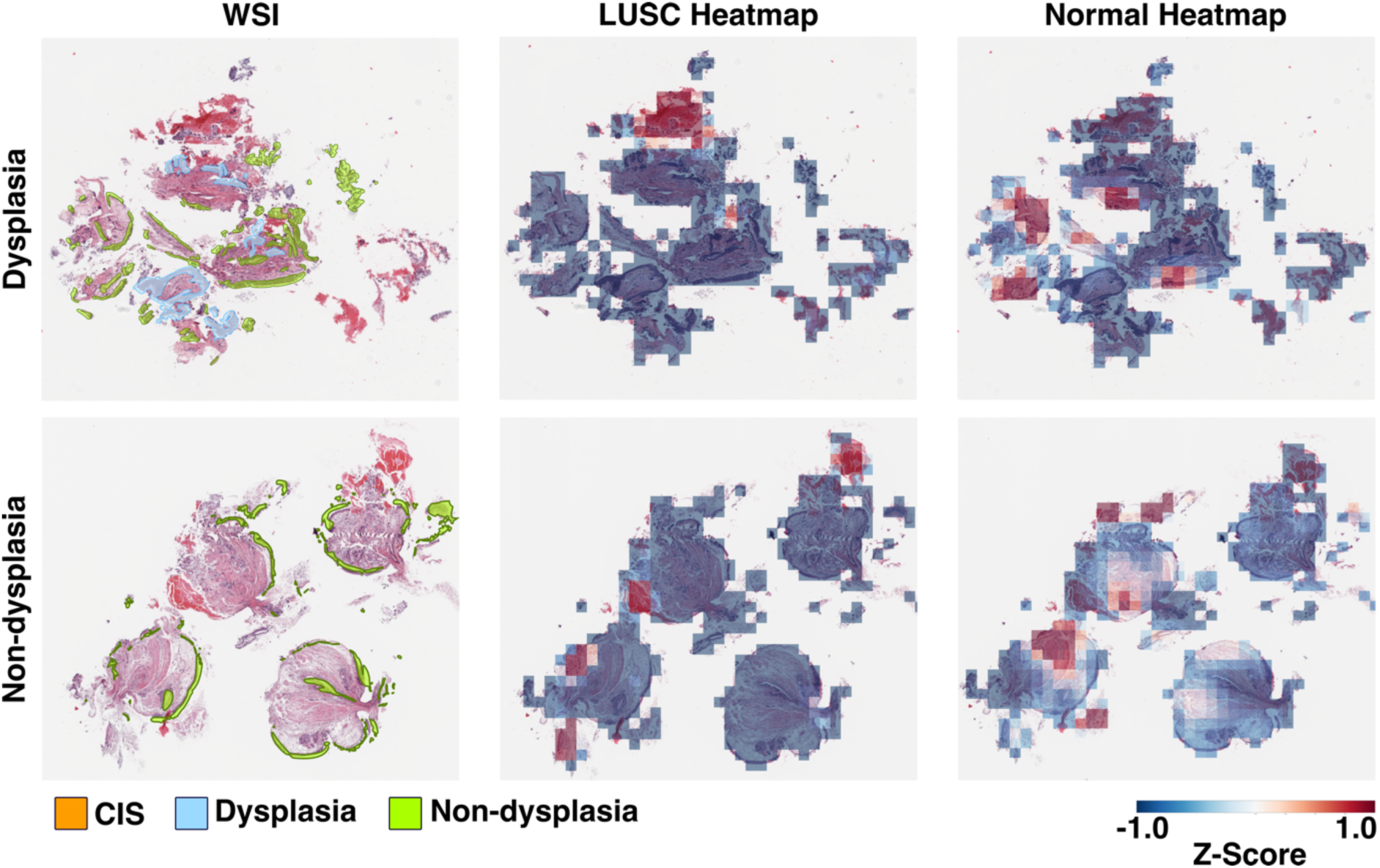
Additional examples of heatmaps generated for Roswell samples: The procedure for explanation visualizations is identical to Fig 2c. WSIs, their overlayed pathologist annotations (1st column) and respective LUSC-specific (2nd column) and normal-specific heatmaps (3rd column) are shown for endobronchial premalignant biopsies from Roswell (top row, a case of with dysplasia regions and bottom row, a case without any regions of dsyplasia) predicted by the GPN to be normal. In cases predicted by the GPN to be normal, the GPN has diculty identifying the small regions of dysplasia with subtle abberrant epithelial changes.

**Supplementary Table 1:** HTAN IDs for the WSIs.

| **FileName** | **HTANBiospecimenID** | **HTANFileID** |
| --- | --- | --- |
| **PCGA-01-0001-029-20126-01119BX.svs** | HTA3_0001_1301119 | HTA3_0001_1901119 |
| **PCGA-01-0001-037-19118-00482BX.svs** | HTA3_0001_1300482 | HTA3_0001_1900482 |
| **PCGA-01-0001-037-19300-00589BX.svs** | HTA3_0001_1300589 | HTA3_0001_1900589 |
| **PCGA-01-0001-037-19762-00898BX.svs** | HTA3_0001_1300898 | HTA3_0001_1900898 |
| **PCGA-01-0001-037-20126-01120BX.svs** | HTA3_0001_1301120 | HTA3_0001_1901120 |
| **PCGA-01-0001-050-15539-90001BX.svs** | HTA3_0001_1390001 | HTA3_0001_1990001 |
| **PCGA-01-0001-050-16918-90004BX.svs** | HTA3_0001_1390004 | HTA3_0001_1990004 |
| **PCGA-01-0001-050-19118-90002BX.svs** | HTA3_0001_1390002 | HTA3_0001_1990002 |
| **PCGA-01-0001-050-19300-90003BX.svs** | HTA3_0001_1390003 | HTA3_0001_1990003 |
| **PCGA-01-0001-050-19762-00903BX.svs** | HTA3_0001_1300903 | HTA3_0001_1900903 |
| **PCGA-01-0001-070-19118-90005BX.svs** | HTA3_0001_1390005 | HTA3_0001_1990005 |
| **PCGA-01-0001-070-19300-00587BX.svs** | HTA3_0001_1300587 | HTA3_0001_1900587 |
| **PCGA-01-0001-070-20126-01117BX.svs** | HTA3_0001_1301117 | HTA3_0001_1901117 |
| **PCGA-01-0001-074-19300-00586BX.svs** | HTA3_0001_1300586 | HTA3_0001_1900586 |
| **PCGA-01-0001-074-20126-01116BX.svs** | HTA3_0001_1301116 | HTA3_0001_1901116 |
| **PCGA-01-0001-074-20343-01315BX.svs** | HTA3_0001_1301315 | HTA3_0001_1901315 |
| **PCGA-01-0001-076-19300-00585BX.svs** | HTA3_0001_1300585 | HTA3_0001_1900585 |
| **PCGA-01-0001-076-20343-01314BX.svs** | HTA3_0001_1301314 | HTA3_0001_1901314 |
| **PCGA-01-0001-077-20126-01115BX.svs** | HTA3_0001_1301115 | HTA3_0001_1901115 |
| **PCGA-01-0001-082-19118-00480BX.svs** | HTA3_0001_1300480 | HTA3_0001_1900480 |
| **PCGA-01-0001-089-19118-00481BX.svs** | HTA3_0001_1300481 | HTA3_0001_1900481 |
| **PCGA-01-0001-089-19300-00588BX.svs** | HTA3_0001_1300588 | HTA3_0001_1900588 |
| **PCGA-01-0001-089-19762-00900BX.svs** | HTA3_0001_1300900 | HTA3_0001_1900900 |
| **PCGA-01-0001-089-20126-01118BX.svs** | HTA3_0001_1301118 | HTA3_0001_1901118 |
| **PCGA-01-0002-021-24062-00441BX.svs** | HTA3_0002_1300441 | HTA3_0002_1900441 |
| **PCGA-01-0002-077-24062-00442BX.svs** | HTA3_0002_1300442 | HTA3_0002_1900442 |
| **PCGA-01-0003-061-23883-90065BX.svs** | HTA3_0003_1390065 | HTA3_0003_1990065 |
| **PCGA-01-0003-061-24282-00694BX.svs** | HTA3_0003_1300694 | HTA3_0003_1900694 |
| **PCGA-01-0003-089-23883-90064BX.svs** | HTA3_0003_1390064 | HTA3_0003_1990064 |
| **PCGA-01-0003-089-24282-00693BX.svs** | HTA3_0003_1300693 | HTA3_0003_1900693 |
| **PCGA-01-0003-089-24730-00947BX.svs** | HTA3_0003_1300947 | HTA3_0003_1900947 |
| **PCGA-01-0004-050-25544-90067BX.svs** | HTA3_0004_1390067 | HTA3_0004_1990067 |
| **PCGA-01-0004-050-25733-90068BX.svs** | HTA3_0004_1390068 | HTA3_0004_1990068 |
| **PCGA-01-0004-050-26477-90006BX.svs** | HTA3_0004_1390006 | HTA3_0004_1990006 |
| **PCGA-01-0004-050-26687-90007BX.svs** | HTA3_0004_1390007 | HTA3_0004_1990007 |
| **PCGA-01-0004-050-27317-00606BX.svs** | HTA3_0004_1300606 | HTA3_0004_1900606 |
| **PCGA-01-0004-089-27317-00604BX.svs** | HTA3_0004_1300604 | HTA3_0004_1900604 |
| **PCGA-01-0004-093-26477-90008BX.svs** | HTA3_0004_1390008 | HTA3_0004_1990008 |
| **PCGA-01-0004-093-26687-90009BX.svs** | HTA3_0004_1390009 | HTA3_0004_1990009 |
| **PCGA-01-0004-093-27317-00605BX.svs** | HTA3_0004_1300605 | HTA3_0004_1900605 |
| **PCGA-01-0005-019-16929-00325BX.svs** | HTA3_0005_1300325 | HTA3_0005_1900325 |
| **PCGA-01-0005-019-17489-00727BX.svs** | HTA3_0005_1300727 | HTA3_0005_1900727 |
| **PCGA-01-0006-037-20238-00776BX.svs** | HTA3_0006_1300776 | HTA3_0006_1900776 |
| **PCGA-01-0006-037-21015-01279BX.svs** | HTA3_0006_1301279 | HTA3_0006_1901279 |
| **PCGA-01-0006-084-19783-00447BX.svs** | HTA3_0006_1300447 | HTA3_0006_1900447 |
| **PCGA-01-0007-048-18033-00455BX.svs** | HTA3_0007_1300455 | HTA3_0007_1900455 |
| **PCGA-01-0007-050-18033-00456BX.svs** | HTA3_0007_1300456 | HTA3_0007_1900456 |
| **PCGA-01-0007-050-18131-00528BX.svs** | HTA3_0007_1300528 | HTA3_0007_1900528 |
| **PCGA-01-0007-089-18033-00453BX.svs** | HTA3_0007_1300453 | HTA3_0007_1900453 |
| **PCGA-01-0007-089-18131-00526BX.svs** | HTA3_0007_1300526 | HTA3_0007_1900526 |
| **PCGA-01-0007-093-18131-00527BX.svs** | HTA3_0007_1300527 | HTA3_0007_1900527 |
| **PCGA-01-0008-045-21682-00840BX.svs** | HTA3_0008_1300840 | HTA3_0008_1900840 |
| **PCGA-01-0008-050-21682-00841BX.svs** | HTA3_0008_1300841 | HTA3_0008_1900841 |
| **PCGA-01-0008-050-22263-01196BX.svs** | HTA3_0008_1301196 | HTA3_0008_1901196 |
| **PCGA-01-0008-074-21682-00837BX.svs** | HTA3_0008_1300837 | HTA3_0008_1900837 |
| **PCGA-01-0008-074-22263-01191BX.svs** | HTA3_0008_1301191 | HTA3_0008_1901191 |
| **PCGA-01-0008-089-21115-00418BX.svs** | HTA3_0008_1300418 | HTA3_0008_1900418 |
| **PCGA-01-0008-089-21682-00838BX.svs** | HTA3_0008_1300838 | HTA3_0008_1900838 |
| **PCGA-01-0008-089-22263-01194BX.svs** | HTA3_0008_1301194 | HTA3_0008_1901194 |
| **PCGA-01-0009-053-23126-00557BX.svs** | HTA3_0009_1300557 | HTA3_0009_1900557 |
| **PCGA-01-0009-053-23497-00844BX.svs** | HTA3_0009_1300844 | HTA3_0009_1900844 |
| **PCGA-01-0009-053-23882-00991BX.svs** | HTA3_0009_1300991 | HTA3_0009_1900991 |
| **PCGA-01-0009-053-24085-01207BX.svs** | HTA3_0009_1301207 | HTA3_0009_1901207 |
| **PCGA-01-0009-061-23126-00558BX.svs** | HTA3_0009_1300558 | HTA3_0009_1900558 |
| **PCGA-01-0009-061-23497-00845BX.svs** | HTA3_0009_1300845 | HTA3_0009_1900845 |
| **PCGA-01-0009-061-23882-00992BX.svs** | HTA3_0009_1300992 | HTA3_0009_1900992 |
| **PCGA-01-0009-092-23497-00846BX.svs** | HTA3_0009_1300846 | HTA3_0009_1900846 |
| **PCGA-01-0009-092-24085-01209BX.svs** | HTA3_0009_1301209 | HTA3_0009_1901209 |
| **PCGA-01-0009-093-21241-90010BX.svs** | HTA3_0009_1390010 | HTA3_0009_1990010 |
| **PCGA-01-0010-048-19599-90011BX.svs** | HTA3_0010_1390011 | HTA3_0010_1990011 |
| **PCGA-01-0010-048-19900-90012BX.svs** | HTA3_0010_1390012 | HTA3_0010_1990012 |
| **PCGA-01-0010-050-19599-00505BX.svs** | HTA3_0010_1300505 | HTA3_0010_1900505 |
| **PCGA-01-0010-050-19599-90013BX.svs** | HTA3_0010_1390013 | HTA3_0010_1990013 |
| **PCGA-01-0010-050-19900-90014BX.svs** | HTA3_0010_1390014 | HTA3_0010_1990014 |
| **PCGA-01-0010-089-19599-00503BX.svs** | HTA3_0010_1300503 | HTA3_0010_1900503 |
| **PCGA-01-0010-089-19599-90061BX.svs** | HTA3_0010_1390061 | HTA3_0010_1990061 |
| **PCGA-01-0010-089-19900-00700BX.svs** | HTA3_0010_1300700 | HTA3_0010_1900700 |
| **PCGA-01-0010-089-19900-90062BX.svs** | HTA3_0010_1390062 | HTA3_0010_1990062 |
| **PCGA-01-0010-094-19599-00504BX.svs** | HTA3_0010_1300504 | HTA3_0010_1900504 |
| **PCGA-01-0010-094-19900-00701BX.svs** | HTA3_0010_1300701 | HTA3_0010_1900701 |
| **PCGA-01-0011-050-26439-00341BX.svs** | HTA3_0011_1300341 | HTA3_0011_1900341 |
| **PCGA-01-0011-050-26558-00433BX.svs** | HTA3_0011_1300433 | HTA3_0011_1900433 |
| **PCGA-01-0011-050-26740-00567BX.svs** | HTA3_0011_1300567 | HTA3_0011_1900567 |
| **PCGA-01-0011-089-26439-00340BX.svs** | HTA3_0011_1300340 | HTA3_0011_1900340 |
| **PCGA-01-0011-089-26558-00432BX.svs** | HTA3_0011_1300432 | HTA3_0011_1900432 |
| **PCGA-01-0011-089-26740-90015BX.svs** | HTA3_0011_1390015 | HTA3_0011_1990015 |
| **PCGA-01-0012-050-21101-00736BX.svs** | HTA3_0012_1300736 | HTA3_0012_1900736 |
| **PCGA-01-0012-050-21836-01253BX.svs** | HTA3_0012_1301253 | HTA3_0012_1901253 |
| **PCGA-01-0014-010-19671-90020BX.svs** | HTA3_0014_1390020 | HTA3_0014_1990020 |
| **PCGA-01-0014-010-20660-00748BX.svs** | HTA3_0014_1300748 | HTA3_0014_1900748 |
| **PCGA-01-0014-011-19323-90017BX.svs** | HTA3_0014_1390017 | HTA3_0014_1990017 |
| **PCGA-01-0014-011-20177-00394BX.svs** | HTA3_0014_1300394 | HTA3_0014_1900394 |
| **PCGA-01-0014-011-20660-00747BX.svs** | HTA3_0014_1300747 | HTA3_0014_1900747 |
| **PCGA-01-0014-011-20660-90018BX.svs** | HTA3_0014_1390018 | HTA3_0014_1990018 |
| **PCGA-01-0014-053-20177-00393BX.svs** | HTA3_0014_1300393 | HTA3_0014_1900393 |
| **PCGA-01-0014-053-20177-90023BX.svs** | HTA3_0014_1390023 | HTA3_0014_1990023 |
| **PCGA-01-0014-053-20660-00745BX.svs** | HTA3_0014_1300745 | HTA3_0014_1900745 |
| **PCGA-01-0014-089-20660-00744BX.svs** | HTA3_0014_1300744 | HTA3_0014_1900744 |
| **PCGA-01-0014-093-20177-90021BX.svs** | HTA3_0014_1390021 | HTA3_0014_1990021 |
| **PCGA-01-0015-011-16303-00402BX.svs** | HTA3_0015_1300402 | HTA3_0015_1900402 |
| **PCGA-01-0015-011-16541-00571BX.svs** | HTA3_0015_1300571 | HTA3_0015_1900571 |
| **PCGA-01-0015-011-17129-00932BX.svs** | HTA3_0015_1300932 | HTA3_0015_1900932 |
| **PCGA-01-0015-011-17638-01327BX.svs** | HTA3_0015_1301327 | HTA3_0015_1901327 |
| **PCGA-01-0015-025-16541-00570BX.svs** | HTA3_0015_1300570 | HTA3_0015_1900570 |
| **PCGA-01-0015-025-16954-00864BX.svs** | HTA3_0015_1300864 | HTA3_0015_1900864 |
| **PCGA-01-0016-025-22821-90024BX.svs** | HTA3_0016_1390024 | HTA3_0016_1990024 |
| **PCGA-01-0016-025-23045-90025BX.svs** | HTA3_0016_1390025 | HTA3_0016_1990025 |
| **PCGA-01-0016-050-22821-00509BX.svs** | HTA3_0016_1300509 | HTA3_0016_1900509 |
| **PCGA-01-0016-050-23045-00647BX.svs** | HTA3_0016_1300647 | HTA3_0016_1900647 |
| **PCGA-01-0017-052-19883-00473BX.svs** | HTA3_0017_1300473 | HTA3_0017_1900473 |
| **PCGA-01-0017-052-19883-90030BX.svs** | HTA3_0017_1390030 | HTA3_0017_1990030 |
| **PCGA-01-0017-052-20261-00732BX.svs** | HTA3_0017_1300732 | HTA3_0017_1900732 |
| **PCGA-01-0017-052-20261-90028BX.svs** | HTA3_0017_1390028 | HTA3_0017_1990028 |
| **PCGA-01-0017-052-20422-00850BX.svs** | HTA3_0017_1300850 | HTA3_0017_1900850 |
| **PCGA-01-0017-052-20422-90029BX.svs** | HTA3_0017_1390029 | HTA3_0017_1990029 |
| **PCGA-01-0017-089-19883-00472BX.svs** | HTA3_0017_1300472 | HTA3_0017_1900472 |
| **PCGA-01-0017-089-20261-00731BX.svs** | HTA3_0017_1300731 | HTA3_0017_1900731 |
| **PCGA-01-0017-089-20422-00849BX.svs** | HTA3_0017_1300849 | HTA3_0017_1900849 |
| **PCGA-01-0018-050-21752-00496BX.svs** | HTA3_0018_1300496 | HTA3_0018_1900496 |
| **PCGA-01-0018-050-21990-00651BX.svs** | HTA3_0018_1300651 | HTA3_0018_1900651 |
| **PCGA-01-0018-050-22732-01146BX.svs** | HTA3_0018_1301146 | HTA3_0018_1901146 |
| **PCGA-01-0018-074-21752-00494BX.svs** | HTA3_0018_1300494 | HTA3_0018_1900494 |
| **PCGA-01-0018-074-21990-00650BX.svs** | HTA3_0018_1300650 | HTA3_0018_1900650 |
| **PCGA-01-0018-074-22732-01145BX.svs** | HTA3_0018_1301145 | HTA3_0018_1901145 |
| **PCGA-01-0020-048-23497-00370BX.svs** | HTA3_0020_1300370 | HTA3_0020_1900370 |
| **PCGA-01-0020-048-23497-90031BX.svs** | HTA3_0020_1390031 | HTA3_0020_1990031 |
| **PCGA-01-0020-048-24050-00784BX.svs** | HTA3_0020_1300784 | HTA3_0020_1900784 |
| **PCGA-01-0020-048-24050-90032BX.svs** | HTA3_0020_1390032 | HTA3_0020_1990032 |
| **PCGA-01-0020-094-23497-00367BX.svs** | HTA3_0020_1300367 | HTA3_0020_1900367 |
| **PCGA-01-0020-094-23497-90033BX.svs** | HTA3_0020_1390033 | HTA3_0020_1990033 |
| **PCGA-01-0020-094-24050-00783BX.svs** | HTA3_0020_1300783 | HTA3_0020_1900783 |
| **PCGA-01-0020-094-24050-90034BX.svs** | HTA3_0020_1390034 | HTA3_0020_1990034 |
| **PCGA-01-0021-010-19826-00723BX.svs** | HTA3_0021_1300723 | HTA3_0021_1900723 |
| **PCGA-01-0021-010-20218-00943BX.svs** | HTA3_0021_1300943 | HTA3_0021_1900943 |
| **PCGA-01-0021-025-19266-00319BX.svs** | HTA3_0021_1300319 | HTA3_0021_1900319 |
| **PCGA-01-0021-025-20218-00942BX.svs** | HTA3_0021_1300942 | HTA3_0021_1900942 |
| **PCGA-01-0021-025-20449-01091BX.svs** | HTA3_0021_1301091 | HTA3_0021_1901091 |
| **PCGA-01-0021-025-20687-01310BX.svs** | HTA3_0021_1301310 | HTA3_0021_1901310 |
| **PCGA-01-0021-050-19434-00465BX.svs** | HTA3_0021_1300465 | HTA3_0021_1900465 |
| **PCGA-01-0021-050-19826-00724BX.svs** | HTA3_0021_1300724 | HTA3_0021_1900724 |
| **PCGA-01-0021-050-20218-00944BX.svs** | HTA3_0021_1300944 | HTA3_0021_1900944 |
| **PCGA-01-0021-050-20449-01092BX.svs** | HTA3_0021_1301092 | HTA3_0021_1901092 |
| **PCGA-01-0021-050-20687-01311BX.svs** | HTA3_0021_1301311 | HTA3_0021_1901311 |
| **PCGA-01-0021-070-19266-90035BX.svs** | HTA3_0021_1390035 | HTA3_0021_1990035 |
| **PCGA-01-0021-070-19623-00581BX.svs** | HTA3_0021_1300581 | HTA3_0021_1900581 |
| **PCGA-01-0021-070-19623-90037BX.svs** | HTA3_0021_1390037 | HTA3_0021_1990037 |
| **PCGA-01-0021-070-19826-00720BX.svs** | HTA3_0021_1300720 | HTA3_0021_1900720 |
| **PCGA-01-0021-070-20218-00940BX.svs** | HTA3_0021_1300940 | HTA3_0021_1900940 |
| **PCGA-01-0021-070-20449-01089BX.svs** | HTA3_0021_1301089 | HTA3_0021_1901089 |
| **PCGA-01-0021-070-20687-01307BX.svs** | HTA3_0021_1301307 | HTA3_0021_1901307 |
| **PCGA-01-0021-088-19826-00721BX.svs** | HTA3_0021_1300721 | HTA3_0021_1900721 |
| **PCGA-01-0021-089-19266-90038BX.svs** | HTA3_0021_1390038 | HTA3_0021_1990038 |
| **PCGA-01-0021-089-19434-90039BX.svs** | HTA3_0021_1390039 | HTA3_0021_1990039 |
| **PCGA-01-0021-089-19623-00582BX.svs** | HTA3_0021_1300582 | HTA3_0021_1900582 |
| **PCGA-01-0021-089-19826-00722BX.svs** | HTA3_0021_1300722 | HTA3_0021_1900722 |
| **PCGA-01-0021-089-19826-90040BX.svs** | HTA3_0021_1390040 | HTA3_0021_1990040 |
| **PCGA-01-0021-089-20218-00941BX.svs** | HTA3_0021_1300941 | HTA3_0021_1900941 |
| **PCGA-01-0021-089-20687-01308BX.svs** | HTA3_0021_1301308 | HTA3_0021_1901308 |
| **PCGA-01-0022-070-20918-00468BX.svs** | HTA3_0022_1300468 | HTA3_0022_1900468 |
| **PCGA-01-0022-070-21548-00876BX.svs** | HTA3_0022_1300876 | HTA3_0022_1900876 |
| **PCGA-01-0022-070-21996-01169BX.svs** | HTA3_0022_1301169 | HTA3_0022_1901169 |
| **PCGA-01-0022-089-20918-00469BX.svs** | HTA3_0022_1300469 | HTA3_0022_1900469 |
| **PCGA-01-0022-089-21996-01170BX.svs** | HTA3_0022_1301170 | HTA3_0022_1901170 |
| **PCGA-01-0022-093-21548-00878BX.svs** | HTA3_0022_1300878 | HTA3_0022_1900878 |
| **PCGA-01-0022-093-21996-01171BX.svs** | HTA3_0022_1301171 | HTA3_0022_1901171 |
| **PCGA-01-0023-010-21429-00830BX.svs** | HTA3_0023_1300830 | HTA3_0023_1900830 |
| **PCGA-01-0023-025-20400-90041BX.svs** | HTA3_0023_1390041 | HTA3_0023_1990041 |
| **PCGA-01-0023-025-20764-90047BX.svs** | HTA3_0023_1390047 | HTA3_0023_1990047 |
| **PCGA-01-0023-026-21429-00827BX.svs** | HTA3_0023_1300827 | HTA3_0023_1900827 |
| **PCGA-01-0023-048-21429-00829BX.svs** | HTA3_0023_1300829 | HTA3_0023_1900829 |
| **PCGA-01-0023-050-20200-90043BX.svs** | HTA3_0023_1390043 | HTA3_0023_1990043 |
| **PCGA-01-0023-050-20246-90042BX.svs** | HTA3_0023_1390042 | HTA3_0023_1990042 |
| **PCGA-01-0023-050-20764-90044BX.svs** | HTA3_0023_1390044 | HTA3_0023_1990044 |
| **PCGA-01-0023-050-21429-00828BX.svs** | HTA3_0023_1300828 | HTA3_0023_1900828 |
| **PCGA-01-0023-061-21429-00826BX.svs** | HTA3_0023_1300826 | HTA3_0023_1900826 |
| **PCGA-01-0023-066-21429-00824BX.svs** | HTA3_0023_1300824 | HTA3_0023_1900824 |
| **PCGA-01-0023-070-20400-90048BX.svs** | HTA3_0023_1390048 | HTA3_0023_1990048 |
| **PCGA-01-0023-070-20764-90045BX.svs** | HTA3_0023_1390045 | HTA3_0023_1990045 |
| **PCGA-01-0023-089-21429-00825BX.svs** | HTA3_0023_1300825 | HTA3_0023_1900825 |
| **PCGA-01-0024-050-23686-00520BX.svs** | HTA3_0024_1300520 | HTA3_0024_1900520 |
| **PCGA-01-0024-050-24050-00780BX.svs** | HTA3_0024_1300780 | HTA3_0024_1900780 |
| **PCGA-01-0025-048-20884-00600BX.svs** | HTA3_0025_1300600 | HTA3_0025_1900600 |
| **PCGA-01-0025-048-21689-01106BX.svs** | HTA3_0025_1301106 | HTA3_0025_1901106 |
| **PCGA-01-0025-050-20576-00384BX.svs** | HTA3_0025_1300384 | HTA3_0025_1900384 |
| **PCGA-01-0025-050-20772-00541BX.svs** | HTA3_0025_1300541 | HTA3_0025_1900541 |
| **PCGA-01-0025-050-20884-00601BX.svs** | HTA3_0025_1300601 | HTA3_0025_1900601 |
| **PCGA-01-0025-050-21164-00807BX.svs** | HTA3_0025_1300807 | HTA3_0025_1900807 |
| **PCGA-01-0025-050-21689-01108BX.svs** | HTA3_0025_1301108 | HTA3_0025_1901108 |
| **PCGA-01-0025-070-20576-90049BX.svs** | HTA3_0025_1390049 | HTA3_0025_1990049 |
| **PCGA-01-0025-070-21164-90050BX.svs** | HTA3_0025_1390050 | HTA3_0025_1990050 |
| **PCGA-01-0025-074-20576-00383BX.svs** | HTA3_0025_1300383 | HTA3_0025_1900383 |
| **PCGA-01-0025-074-20772-00540BX.svs** | HTA3_0025_1300540 | HTA3_0025_1900540 |
| **PCGA-01-0025-074-20884-00599BX.svs** | HTA3_0025_1300599 | HTA3_0025_1900599 |
| **PCGA-01-0025-074-21164-00805BX.svs** | HTA3_0025_1300805 | HTA3_0025_1900805 |
| **PCGA-01-0025-074-21689-01105BX.svs** | HTA3_0025_1301105 | HTA3_0025_1901105 |
| **PCGA-01-0025-094-21164-00806BX.svs** | HTA3_0025_1300806 | HTA3_0025_1900806 |
| **PCGA-01-0025-094-21689-01107BX.svs** | HTA3_0025_1301107 | HTA3_0025_1901107 |
| **PCGA-01-0026-089-21752-00337BX.svs** | HTA3_0026_1300337 | HTA3_0026_1900337 |
| **PCGA-01-0026-089-22473-00859BX.svs** | HTA3_0026_1300859 | HTA3_0026_1900859 |
| **PCGA-01-0027-089-19515-00639BX.svs** | HTA3_0027_1300639 | HTA3_0027_1900639 |
| **PCGA-01-0028-041-23527-00757BX.svs** | HTA3_0028_1300757 | HTA3_0028_1900757 |
| **PCGA-01-0028-041-24262-01264BX.svs** | HTA3_0028_1301264 | HTA3_0028_1901264 |
| **PCGA-01-0028-050-23163-00491BX.svs** | HTA3_0028_1300491 | HTA3_0028_1900491 |
| **PCGA-01-0028-050-23527-00758BX.svs** | HTA3_0028_1300758 | HTA3_0028_1900758 |
| **PCGA-01-0028-050-24262-01265BX.svs** | HTA3_0028_1301265 | HTA3_0028_1901265 |
| **PCGA-01-0029-045-22412-00625BX.svs** | HTA3_0029_1300625 | HTA3_0029_1900625 |
| **PCGA-01-0029-045-22678-00814BX.svs** | HTA3_0029_1300814 | HTA3_0029_1900814 |
| **PCGA-01-0029-048-22090-00379BX.svs** | HTA3_0029_1300379 | HTA3_0029_1900379 |
| **PCGA-01-0029-048-22412-90051BX.svs** | HTA3_0029_1390051 | HTA3_0029_1990051 |
| **PCGA-01-0029-048-22678-00815BX.svs** | HTA3_0029_1300815 | HTA3_0029_1900815 |
| **PCGA-01-0029-050-22090-00380BX.svs** | HTA3_0029_1300380 | HTA3_0029_1900380 |
| **PCGA-01-0029-050-22678-00816BX.svs** | HTA3_0029_1300816 | HTA3_0029_1900816 |
| **PCGA-01-0029-061-22090-00378BX.svs** | HTA3_0029_1300378 | HTA3_0029_1900378 |
| **PCGA-01-0029-061-22412-00623BX.svs** | HTA3_0029_1300623 | HTA3_0029_1900623 |
| **PCGA-01-0029-061-22678-00812BX.svs** | HTA3_0029_1300812 | HTA3_0029_1900812 |
| **PCGA-01-0029-070-22090-00376BX.svs** | HTA3_0029_1300376 | HTA3_0029_1900376 |
| **PCGA-01-0029-070-22412-90052BX.svs** | HTA3_0029_1390052 | HTA3_0029_1990052 |
| **PCGA-01-0029-070-22678-00810BX.svs** | HTA3_0029_1300810 | HTA3_0029_1900810 |
| **PCGA-01-0029-089-22090-00377BX.svs** | HTA3_0029_1300377 | HTA3_0029_1900377 |
| **PCGA-01-0029-089-22412-00622BX.svs** | HTA3_0029_1300622 | HTA3_0029_1900622 |
| **PCGA-01-0029-093-22090-90054BX.svs** | HTA3_0029_1390054 | HTA3_0029_1990054 |
| **PCGA-01-0029-093-22412-00624BX.svs** | HTA3_0029_1300624 | HTA3_0029_1900624 |
| **PCGA-01-0029-093-22678-00813BX.svs** | HTA3_0029_1300813 | HTA3_0029_1900813 |
| **PCGA-01-0030-089-27868-00358BX.svs** | HTA3_0030_1300358 | HTA3_0030_1900358 |
| **PCGA-01-0031-021-24061-00664BX.svs** | HTA3_0031_1300664 | HTA3_0031_1900664 |
| **PCGA-01-0031-021-24726-01062BX.svs** | HTA3_0031_1301062 | HTA3_0031_1901062 |
| **PCGA-01-0031-050-23830-00537BX.svs** | HTA3_0031_1300537 | HTA3_0031_1900537 |
| **PCGA-01-0031-050-24061-00666BX.svs** | HTA3_0031_1300666 | HTA3_0031_1900666 |
| **PCGA-01-0031-050-24285-00856BX.svs** | HTA3_0031_1300856 | HTA3_0031_1900856 |
| **PCGA-01-0031-050-24726-01063BX.svs** | HTA3_0031_1301063 | HTA3_0031_1901063 |
| **PCGA-01-0031-053-24061-00665BX.svs** | HTA3_0031_1300665 | HTA3_0031_1900665 |
| **PCGA-01-0031-053-24285-00855BX.svs** | HTA3_0031_1300855 | HTA3_0031_1900855 |
| **PCGA-01-0031-053-24285-90057BX.svs** | HTA3_0031_1390057 | HTA3_0031_1990057 |
| **PCGA-01-0031-053-24726-01061BX.svs** | HTA3_0031_1301061 | HTA3_0031_1901061 |
| **PCGA-01-0031-066-23830-90055BX.svs** | HTA3_0031_1390055 | HTA3_0031_1990055 |
| **PCGA-01-0031-066-24061-90056BX.svs** | HTA3_0031_1390056 | HTA3_0031_1990056 |
| **PCGA-01-0031-074-24285-00854BX.svs** | HTA3_0031_1300854 | HTA3_0031_1900854 |
| **PCGA-01-0031-074-24726-01060BX.svs** | HTA3_0031_1301060 | HTA3_0031_1901060 |
| **PCGA-01-0032-088-24051-00544BX.svs** | HTA3_0032_1300544 | HTA3_0032_1900544 |
| **PCGA-01-0033-048-20582-00974BX.svs** | HTA3_0033_1300974 | HTA3_0033_1900974 |
| **PCGA-01-0033-048-20864-01248BX.svs** | HTA3_0033_1301248 | HTA3_0033_1901248 |
| **PCGA-01-0033-048-21079-01371BX.svs** | HTA3_0033_1301371 | HTA3_0033_1901371 |
| **PCGA-01-0033-050-20582-00977BX.svs** | HTA3_0033_1300977 | HTA3_0033_1900977 |
| **PCGA-01-0033-050-20864-01249BX.svs** | HTA3_0033_1301249 | HTA3_0033_1901249 |
| **PCGA-01-0033-061-20864-01246BX.svs** | HTA3_0033_1301246 | HTA3_0033_1901246 |
| **PCGA-01-0033-085-20864-01245BX.svs** | HTA3_0033_1301245 | HTA3_0033_1901245 |
| **PCGA-01-0033-085-21079-01370BX.svs** | HTA3_0033_1301370 | HTA3_0033_1901370 |
| **PCGA-01-0033-092-21079-01369BX.svs** | HTA3_0033_1301369 | HTA3_0033_1901369 |
| **PCGA-01-0034-050-22969-00918BX.svs** | HTA3_0034_1300918 | HTA3_0034_1900918 |
| **PCGA-01-0035-019-21168-01034BX.svs** | HTA3_0035_1301034 | HTA3_0035_1901034 |
| **PCGA-01-0035-019-21364-01259BX.svs** | HTA3_0035_1301259 | HTA3_0035_1901259 |
| **PCGA-01-0035-042-21168-01035BX.svs** | HTA3_0035_1301035 | HTA3_0035_1901035 |
| **PCGA-01-0035-042-21364-01260BX.svs** | HTA3_0035_1301260 | HTA3_0035_1901260 |
| **PCGA-01-0035-050-21168-01039BX.svs** | HTA3_0035_1301039 | HTA3_0035_1901039 |
| **PCGA-01-0035-061-21168-01037BX.svs** | HTA3_0035_1301037 | HTA3_0035_1901037 |
| **PCGA-01-0035-061-21364-01257BX.svs** | HTA3_0035_1301257 | HTA3_0035_1901257 |
| **PCGA-01-0035-074-21168-01036BX.svs** | HTA3_0035_1301036 | HTA3_0035_1901036 |
| **PCGA-01-0035-074-21364-01256BX.svs** | HTA3_0035_1301256 | HTA3_0035_1901256 |
| **PCGA-01-0035-093-21168-01038BX.svs** | HTA3_0035_1301038 | HTA3_0035_1901038 |
| **PCGA-01-0035-093-21364-01258BX.svs** | HTA3_0035_1301258 | HTA3_0035_1901258 |
| **PCGA-01-0036-048-21975-00298BX.svs** | HTA3_0036_1300298 | HTA3_0036_1900298 |
| **PCGA-01-0036-048-22885-00913BX.svs** | HTA3_0036_1300913 | HTA3_0036_1900913 |
| **PCGA-01-0036-050-22885-00914BX.svs** | HTA3_0036_1300914 | HTA3_0036_1900914 |
| **PCGA-01-0036-050-23647-01405BX.svs** | HTA3_0036_1301405 | HTA3_0036_1901405 |
| **PCGA-01-0036-089-21975-00297BX.svs** | HTA3_0036_1300297 | HTA3_0036_1900297 |
| **PCGA-01-0036-089-22885-00911BX.svs** | HTA3_0036_1300911 | HTA3_0036_1900911 |
| **PCGA-01-0036-089-23647-01404BX.svs** | HTA3_0036_1301404 | HTA3_0036_1901404 |
| **PCGA-01-0037-089-27849-00344BX.svs** | HTA3_0037_1300344 | HTA3_0037_1900344 |
| **PCGA-01-0037-089-28409-00751BX.svs** | HTA3_0037_1300751 | HTA3_0037_1900751 |
| **PCGA-01-0038-037-20756-00770BX.svs** | HTA3_0038_1300770 | HTA3_0038_1900770 |
| **PCGA-01-0038-037-21323-01075BX.svs** | HTA3_0038_1301075 | HTA3_0038_1901075 |
| **PCGA-01-0038-037-21699-01390BX.svs** | HTA3_0038_1301390 | HTA3_0038_1901390 |
| **PCGA-01-0038-048-21323-01076BX.svs** | HTA3_0038_1301076 | HTA3_0038_1901076 |
| **PCGA-01-0038-048-21699-01391BX.svs** | HTA3_0038_1301391 | HTA3_0038_1901391 |
| **PCGA-01-0038-050-20756-00771BX.svs** | HTA3_0038_1300771 | HTA3_0038_1900771 |
| **PCGA-01-0038-050-21323-01077BX.svs** | HTA3_0038_1301077 | HTA3_0038_1901077 |
| **PCGA-01-0038-050-21699-01392BX.svs** | HTA3_0038_1301392 | HTA3_0038_1901392 |
| **PCGA-01-0039-066-19554-01003BX.svs** | HTA3_0039_1301003 | HTA3_0039_1901003 |
| **PCGA-01-0039-076-19554-01002BX.svs** | HTA3_0039_1301002 | HTA3_0039_1901002 |
| **PCGA-01-0039-076-19757-01220BX.svs** | HTA3_0039_1301220 | HTA3_0039_1901220 |
| **PCGA-01-0039-089-19554-01004BX.svs** | HTA3_0039_1301004 | HTA3_0039_1901004 |
| **PCGA-01-0039-089-19757-01222BX.svs** | HTA3_0039_1301222 | HTA3_0039_1901222 |
| **PCGA-01-0039-093-19554-01005BX.svs** | HTA3_0039_1301005 | HTA3_0039_1901005 |
| **PCGA-01-0039-093-19757-01224BX.svs** | HTA3_0039_1301224 | HTA3_0039_1901224 |
| **PCGA-01-0040-050-19706-01100BX.svs** | HTA3_0040_1301100 | HTA3_0040_1901100 |
| **PCGA-01-0040-050-19895-01284BX.svs** | HTA3_0040_1301284 | HTA3_0040_1901284 |
| **PCGA-01-0040-093-19706-01098BX.svs** | HTA3_0040_1301098 | HTA3_0040_1901098 |
| **PCGA-01-0040-093-19895-01283BX.svs** | HTA3_0040_1301283 | HTA3_0040_1901283 |
| **PCGA-01-0041-050-21430-00796BX.svs** | HTA3_0041_1300796 | HTA3_0041_1900796 |
| **PCGA-01-0041-050-21703-00922BX.svs** | HTA3_0041_1300922 | HTA3_0041_1900922 |
| **PCGA-01-0041-050-22044-01183BX.svs** | HTA3_0041_1301183 | HTA3_0041_1901183 |
| **PCGA-01-0041-093-21430-00795BX.svs** | HTA3_0041_1300795 | HTA3_0041_1900795 |
| **PCGA-01-0041-093-21703-00919BX.svs** | HTA3_0041_1300919 | HTA3_0041_1900919 |
| **PCGA-01-0041-093-22044-01182BX.svs** | HTA3_0041_1301182 | HTA3_0041_1901182 |
| **PCGA-01-0042-010-23680-00906BX.svs** | HTA3_0042_1300906 | HTA3_0042_1900906 |
| **PCGA-01-0042-010-23911-01028BX.svs** | HTA3_0042_1301028 | HTA3_0042_1901028 |
| **PCGA-01-0042-048-23680-00907BX.svs** | HTA3_0042_1300907 | HTA3_0042_1900907 |
| **PCGA-01-0042-048-23911-01029BX.svs** | HTA3_0042_1301029 | HTA3_0042_1901029 |
| **PCGA-01-0043-037-21235-01043BX.svs** | HTA3_0043_1301043 | HTA3_0043_1901043 |
| **PCGA-01-0043-037-21599-01350BX.svs** | HTA3_0043_1301350 | HTA3_0043_1901350 |
| **PCGA-01-0043-085-21599-01349BX.svs** | HTA3_0043_1301349 | HTA3_0043_1901349 |
| **PCGA-01-0044-048-17646-44967BX.svs** | HTA3_0044_1300967 | HTA3_0044_1900967 |
| **PCGA-01-0044-048-17898-01132BX.svs** | HTA3_0044_1301132 | HTA3_0044_1901132 |
| **PCGA-01-0044-050-17646-44968BX.svs** | HTA3_0044_1300968 | HTA3_0044_1900968 |
| **PCGA-01-0044-050-17898-01133BX.svs** | HTA3_0044_1301133 | HTA3_0044_1901133 |
| **PCGA-01-0044-050-18295-01397BX.svs** | HTA3_0044_1301397 | HTA3_0044_1901397 |
| **PCGA-01-0044-070-16659-00293BX.svs** | HTA3_0044_1300293 | HTA3_0044_1900293 |
| **PCGA-01-0044-070-17646-44965BX.svs** | HTA3_0044_1300965 | HTA3_0044_1900965 |
| **PCGA-01-0044-070-17898-01130BX.svs** | HTA3_0044_1301130 | HTA3_0044_1901130 |
| **PCGA-01-0044-089-17646-44966BX.svs** | HTA3_0044_1300966 | HTA3_0044_1900966 |
| **PCGA-01-0044-089-17898-01131BX.svs** | HTA3_0044_1301131 | HTA3_0044_1901131 |
| **PCGA-01-0044-089-18295-01396BX.svs** | HTA3_0044_1301396 | HTA3_0044_1901396 |
| **PCGA-01-0045-045-17021-00893BX.svs** | HTA3_0045_1300893 | HTA3_0045_1900893 |
| **PCGA-01-0045-048-17021-00894BX.svs** | HTA3_0045_1300894 | HTA3_0045_1900894 |
| **PCGA-01-0045-048-17497-01241BX.svs** | HTA3_0045_1301241 | HTA3_0045_1901241 |
| **PCGA-01-0045-085-17021-00891BX.svs** | HTA3_0045_1300891 | HTA3_0045_1900891 |
| **PCGA-01-0045-085-17497-01238BX.svs** | HTA3_0045_1301238 | HTA3_0045_1901238 |
| **PCGA-01-0046-047-26739-00766BX.svs** | HTA3_0046_1300766 | HTA3_0046_1900766 |
| **PCGA-01-0046-047-27551-01292BX.svs** | HTA3_0046_1301292 | HTA3_0046_1901292 |
| **PCGA-01-0046-050-27138-00953BX.svs** | HTA3_0046_1300953 | HTA3_0046_1900953 |
| **PCGA-01-0046-050-27551-01293BX.svs** | HTA3_0046_1301293 | HTA3_0046_1901293 |
| **PCGA-01-0046-089-26739-00765BX.svs** | HTA3_0046_1300765 | HTA3_0046_1900765 |
| **PCGA-01-0046-089-27138-00952BX.svs** | HTA3_0046_1300952 | HTA3_0046_1900952 |
| **PCGA-01-0047-037-19263-01021BX.svs** | HTA3_0047_1301021 | HTA3_0047_1901021 |
| **PCGA-01-0047-037-19447-01216BX.svs** | HTA3_0047_1301216 | HTA3_0047_1901216 |
| **PCGA-01-0047-037-19809-01411BX.svs** | HTA3_0047_1301411 | HTA3_0047_1901411 |
| **PCGA-01-0047-050-19263-01022BX.svs** | HTA3_0047_1301022 | HTA3_0047_1901022 |
| **PCGA-01-0048-025-25305-48965BX.svs** | HTA3_0048_1300965 | HTA3_0048_1900965 |
| **PCGA-01-0048-025-25669-01275BX.svs** | HTA3_0048_1301275 | HTA3_0048_1901275 |
| **PCGA-01-0048-061-25669-01273BX.svs** | HTA3_0048_1301273 | HTA3_0048_1901273 |
| **PCGA-01-0048-061-25835-01381BX.svs** | HTA3_0048_1301381 | HTA3_0048_1901381 |
| **PCGA-01-0048-093-25102-00882BX.svs** | HTA3_0048_1300882 | HTA3_0048_1900882 |
| **PCGA-01-0048-093-25835-01380BX.svs** | HTA3_0048_1301380 | HTA3_0048_1901380 |
| **PCGA-01-0049-010-23043-01187BX.svs** | HTA3_0049_1301187 | HTA3_0049_1901187 |
| **PCGA-01-0050-050-23482-00998BX.svs** | HTA3_0050_1300998 | HTA3_0050_1900998 |
| **PCGA-01-0050-050-23699-01235BX.svs** | HTA3_0050_1301235 | HTA3_0050_1901235 |
| **PCGA-01-0050-084-23482-00997BX.svs** | HTA3_0050_1300997 | HTA3_0050_1900997 |
| **PCGA-01-0051-089-22577-01162BX.svs** | HTA3_0051_1301162 | HTA3_0051_1901162 |
| **PCGA-01-0051-089-22786-01334BX.svs** | HTA3_0051_1301334 | HTA3_0051_1901334 |
| **PCGA-01-0052-036-20842-01018BX.svs** | HTA3_0052_1301018 | HTA3_0052_1901018 |
| **PCGA-01-0052-036-21157-01301BX.svs** | HTA3_0052_1301301 | HTA3_0052_1901301 |
| **PCGA-01-0052-048-20842-01017BX.svs** | HTA3_0052_1301017 | HTA3_0052_1901017 |
| **PCGA-01-0052-048-21157-01302BX.svs** | HTA3_0052_1301302 | HTA3_0052_1901302 |
| **PCGA-01-0052-050-20842-01015BX.svs** | HTA3_0052_1301015 | HTA3_0052_1901015 |
| **PCGA-01-0052-093-20842-01014BX.svs** | HTA3_0052_1301014 | HTA3_0052_1901014 |
| **PCGA-01-0052-093-21157-01299BX.svs** | HTA3_0052_1301299 | HTA3_0052_1901299 |

